# Mechanism of broad-spectrum Cas9 inhibition by AcrIIA11

**DOI:** 10.1101/2021.09.15.460536

**Authors:** Kaylee E. Dillard, Cynthia Terrace, Kamyab Javanmardi, Wantae Kim, Kevin J. Forsberg, Ilya J. Finkelstein

**Affiliations:** Department of Molecular Biosciences and Institute for Cellular and Molecular Biology, University of Texas at Austin, Austin, Texas 78712, USA; Center for Systems and Synthetic Biology, University of Texas at Austin, Austin, Texas 78712, USA; Department of Microbiology, University of Texas Southwestern Medical Center, Dallas, Texas, 75390, USA

**Keywords:** Anti-CRISPR, single-molecule, DNA Curtains, Acr

## Abstract

Mobile genetic elements evade CRISPR-Cas adaptive immunity by encoding anti-CRISPR proteins (Acrs). Acrs inactivate CRISPR-Cas systems via diverse mechanisms but are generally specific for a narrow subset of Cas nucleases that share high sequence similarity. Here, we demonstrate that AcrIIA11 inhibits diverse Cas9 sub-types *in vitro* and human cells. Single-molecule fluorescence imaging reveals that AcrIIA11 interferes with the first steps of target search by reducing *S. aureus* Cas9’s diffusion on non-specific DNA. DNA cleavage is inhibited because the AcrIIA11:Cas9 complex is kinetically trapped at PAM-rich decoy sites, preventing Cas9 from reaching its target. This work establishes that DNA trapping can be used to inhibit a broad spectrum of Cas9 orthologs *in vitro* and during mammalian genome editing.

## Introduction

The molecular arms race between prokaryotes and mobile genetic elements (MGEs) has resulted in proteins that suppress CRISPR-Cas adaptive immunity. CRISPR-Cas systems protect bacteria and archaea from MGEs by incorporating a fragment of the foreign nucleic acid into the host genome as a spacer. This spacer is transcribed into a CRISPR RNA (crRNA) and assembled with CRISPR-Cas proteins into an effector complex. The effector complex can then target and degrade the MGEs upon reinfection. Targeting proceeds via a multi-step mechanism (1–4). First, the effector complex binds a protospacer-adjacent motif (PAM) via protein-DNA interactions. At a strong PAM, the duplex DNA partially unwinds and pairs with the crRNA to form an RNA:DNA hybrid (R-loop). Nucleolytic degradation of the target DNA occurs after stable R-loop formation (1, 4–7). However, MGEs have developed diverse ways to circumvent CRISPR-Cas adaptive immunity. For example, phages and prophages can encode anti-CRISPR proteins (Acrs) that interfere with crRNA and CRISPR effector complex biogenesis, DNA binding, and nuclease activation (8–10). The first Acrs were discovered in *Pseudomonas aeruginosa* lysogens containing an active Type I-F CRISPR-Cas system that was sensitive to phage plaquing (11–16). Since this pioneering work, over 50 putative Acrs have been discovered via bioinformatics, molecular biology, and functional metagenomic approaches (9, 16–23). The mechanism of action of some of these Acrs remains unknown.

Acrs that inactivate Class 2 Cas nucleases have received increasing attention due to their potential biotechnological applications (17, 18, 24–26). For example, AcrIIA4, a DNA-mimicking inhibitor of *S. pyogenes (Sp)*Cas9, has been used to selectively deactivate gene editing in some tissues (27, 28), promote homology-directed repair by limiting DNA cleavage to S/G2 phases (29), and regulate CRISPRa (CRISPR gene activation)/CRISPRi (CRISPR gene interference) in synthetic gene circuits (30). We previously discovered AcrIIA11 as a potent *Sp*Cas9 inhibitor via a high-throughput functional metagenomic screen (22). We showed that AcrIIA11 inhibited *Sp*Cas9 *in vitro*, in *E*.*coli*, and mammalian cells via an unknown mechanism that appeared distinct from any Class 2 Acr mechanism to date. Our earlier study also did not explore whether AcrIIA11 could inhibit other Cas9 orthologs.

Here, we show that AcrIIA11 is a broad-spectrum inhibitor of multiple Cas9 orthologs both *in vitro* and in human cells. Biochemical experiments indicate that AcrIIA11 binds and strongly inhibits *Sa*Cas9 ribonucleoprotein (RNP) (compared to *Sp*Cas9, *Nme*Cas9, and *Fn*Cas9). Using single-molecule imaging, we demonstrate that *Sa*Cas9 RNPs slide on double-stranded DNA (dsDNA) in search of the crRNA-complementary target DNA. However, AcrIIA11:*Sa*Cas9 complexes restrict this one-dimensional diffusion on DNA and retain the nuclease at PAM-rich off-target decoy sites. By tethering *Sa*Cas9 to PAM-rich off-target sites, AcrIIA11 blocks Cas9’s ability to find and recognize its target. Collectively, our results indicate that AcrIIA11 inhibits diverse Cas9 orthologs by capturing the enzymes on off-target DNA sites.

## Results

### AcrIIA11 inhibits Cas9 orthologs in vitro and in human cells

To better understand the mechanisms of Cas9 inhibition, we assayed whether AcrIIA11 can inhibit DNA cleavage by *Sa*Cas9 (Type II-A), *Fn*Cas9 (Type II-B), and *Nme*Cas9 (Type II-C). AcrIIA11 was incubated with each Cas9 ortholog before adding single guide RNA (sgRNA) and a linearized plasmid containing the target DNA adjacent to the Cas9-specific PAM. Cleavage was assayed at various timepoints and resolved on an agarose gel (**Figure 1**). Consistent with a prior report (40), we extended the incubation time for *Fn*Cas9, which binds and cleaves DNA much slower than *Sa*Cas9 or *Nme*Cas9. AcrIIA11 efficiently inhibited *Sa*Cas9 and *Fn*Cas9 but did not inhibit *Nme*Cas9 (**Figures 1E-F**). We also ruled out that AcrIIA11 harbors RNAse activity on the sgRNAs (**Figure S1**). *Sa*Cas9 inhibition was more rapid and more complete than what we observed with all other Cas9 variants (22). Since *Sa*Cas9 is inhibited strongly and rapidly by AcrIIA11 *in vitro*, and also because it is frequently used for mammalian gene editing, we selected this enzyme for all further experiments (41, 42).

**Figure 1.**
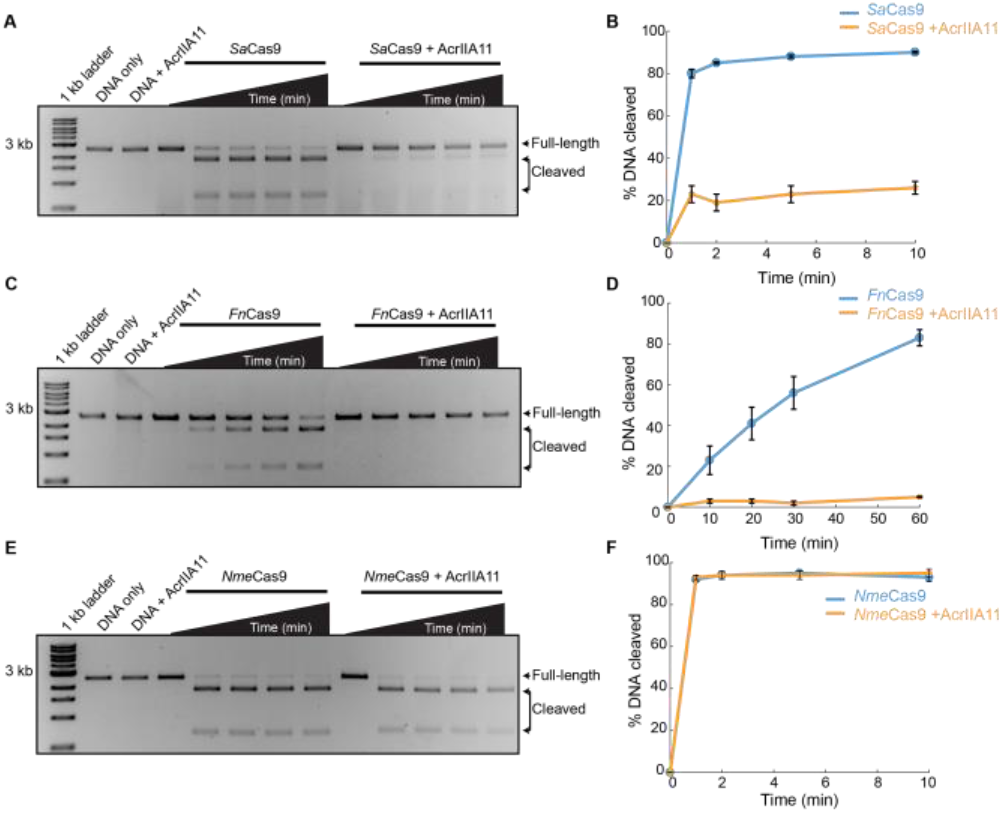
AcrIIA11 inhibits Cas9 orthologs *in vitro*. Agarose gels and quantification of DNA cleavage with (A, B) *Sa*Cas9, (C, D) *Fn*Cas9, and (E, F) *Nme*Cas9. Time points for *Sa*Cas9 and *Nme*Cas9: 0, 1, 2, 5, 10 min. Time points for *Fn*Cas9: 0, 10, 20, 30, 60 min. Graphs represent the mean of three replicates. Error bars: S.E.M.

Next, we tested whether AcrIIA11 can inhibit *Sa*Cas9 in human cell cultures. HEK293Ts were co-transfected with two plasmids. The first plasmid expressed *Sa*Cas9 along with a sgRNA; the second expressed AcrIIA11 or AcrIIA4 (**Figure 2 and Figure S2**). AcrIIA4 was included as a control because it inhibits *Sp*Cas9 but not *Sa*Cas9 (43). As expected, AcrIIA4 did not inhibit *Sa*Cas9 at any of the four tested loci. We also did not see any cleavage when a scrambled sgRNA was used in the absence of any inhibitors. AcrIIA11 inhibited *Sa*Cas9 at the *CACNA1D, EMX1*, and *FANCF* loci (**Figure 2 and S2A-D**). Surprisingly, AcrIIA11 did not inhibit *Sa*Cas9 cleavage at a fourth site, *RUNX1* (**Figure 2D and S2E-F**). The overall indel percentage at *RUNX1* was higher than all other targets, suggesting that this site is more accessible to *Sa*Cas9. We previously showed that AcrIIA11 also inhibits *Sp*Cas9 at *CACNA1D*, but not at *EMX1* (22). Together, these data indicate AcrIIA11 is a robust inhibitor of Cas9 orthologs *in vitro* and a locus-specific inhibitor in human cells.

**Figure 2.**
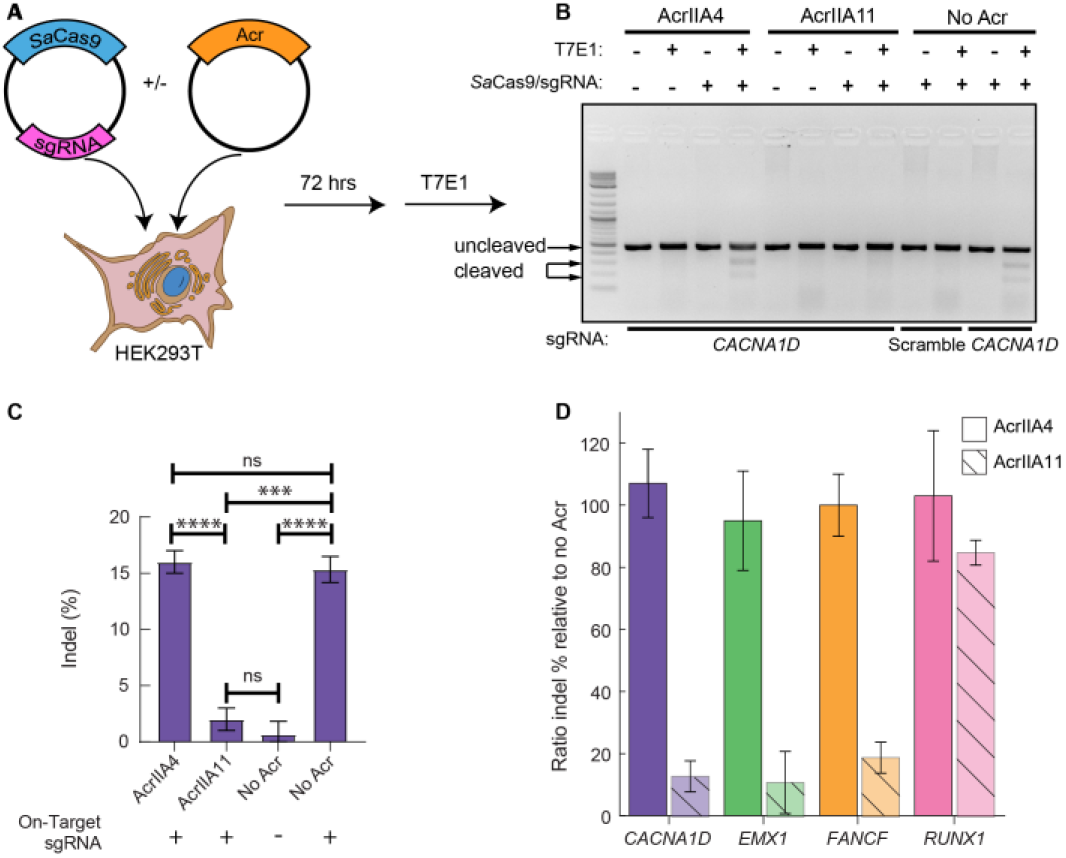
AcrIIA11 inhibits *Sa*Cas9 in human cells. (A) HEK293T cells are transiently transfected with plasmids carrying *Sa*Cas9 + sgRNA. A second plasmid encoding either AcrIIA4 or AcrIIA11 is included, as indicated. (B, C) Representative agarose gel and quantification of indel percentage for the *CACNA1D* site. Error bars are the standard deviation of three replicates. P-values (not significant [ns], p > 0.05; *p < 0.05; **p < 0.01; ***p < 0.001; ****p < 0.0001) were determined using a Student’s t-test. (D) Quantification of the editing efficiency when AcrIIA11 or AcrIIA4 are present relative to when no Acr is expressed.

### AcrIIA11 sequesters Cas9 at off-target sites to prevent target recognition

Many Acrs directly bind their CRISPR effector complex, and we hypothesized that AcrIIA11 may also bind Cas9 (12, 14, 44–54). Previously, we showed that AcrIIA11 interacts with *Sp*Cas9 with a slight preference for the *Sp*Cas9-sgRNA RNP (22). Here, we confirmed that AcrIIA11 also binds *Sa*Cas9. For these experiments, the *Sa*Cas9 RNP was purified with an N-terminal TwinStrep-SUMO epitope and immobilized on a Strep-Tactin resin. We could readily capture AcrIIA11 on the resin-immobilized *Sa*Cas9, and *Sa*Cas9 (lacking the TwinStrep epitope) was captured when we immobilized AcrIIA11 (**Figure S3A, C**). The AcrIIA11:*Sa*Cas9 complex was eluted from the Strep-Tactin resin and further purified on a size exclusion column. Since AcrIIA11 can weakly bind sgRNA (**Figure S1**), we tested if apo*Sa*Cas9 (lacking sgRNA) could capture AcrIIA11. Immobilized apo*Sa*Cas9 captured AcrIIA11, and immobilized AcrIIA11 could also capture apo*Sa*Cas9 (**Figure S3B, D**) We conclude that AcrIIA11 binds Cas9s via protein-protein interactions, and these interactions may be stabilized by AcrIIA11-sgRNA contacts.

To understand how AcrIIA11 inhibits Cas9, we visualized individual *Sa*Cas9 molecules as they scan dsDNA for their target sites (**Figure 3A**). In this assay, a 48.5 kb-long DNA substrate is suspended above a lipid bilayer between two microfabricated chromium features (37, 38). The DNA is prepared with a biotin on one end and a digoxigenin on the opposite end. The biotinylated end is immobilized on the surface of a fluid lipid bilayer via a biotin-streptavidin linkage. In addition to capturing one end of the DNA substrate, the bilayer also passivates the flowcell surface for biochemical experiments. Next, the DNA molecules are organized and extended at microfabricated chrome barriers via buffer flow. The second DNA end is captured at the anti-digoxigenin-functionalized chrome pedestal and buffer flow is terminated. Our attempts to fluorescently label AcrIIA11 resulted in a partial loss of activity. Therefore, we fluorescently labeled *Sa*Cas9 via a fluorescent anti-FLAG antibody that targeted a 3xFLAG epitope on its N-terminus. Using dual-tethered DNA molecules, we can thus track *Sa*Cas9 diffusion in the absence of buffer flow.

**Figure 3.**
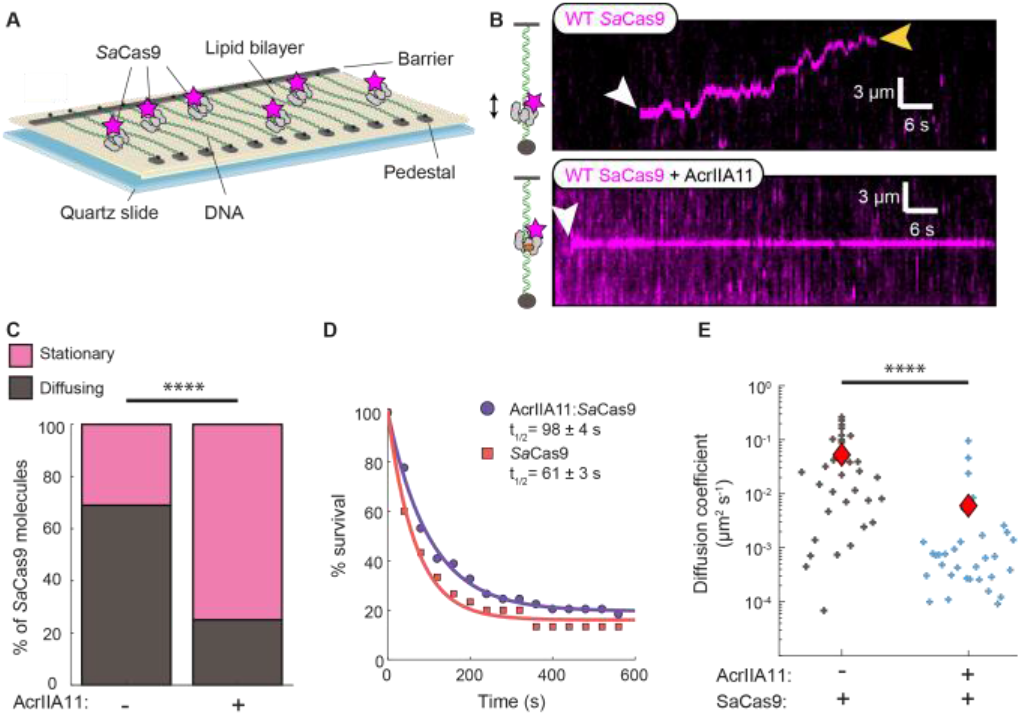
AcrIIA11 retains *Sa*Cas9 on non-specific DNA to slow the target search dynamics. (A) Schematic of the double-tethered DNA curtains assay. Buffer flow is turned off after both ends of the DNA are tethered between the chromium barriers and pedestals and the protein enters the flowcell. (B) Kymographs showing diffusing *Sa*Cas9 RNPs (top) and stationary AcrIIA11:*Sa*Cas9 RNP complexes (bottom). The white arrow indicates the time when *Sa*Cas9 binds, and the yellow arrow indicates when *Sa*Cas9 releases the DNA (C) Most *Sa*Cas9 RNPs diffuse without AcrIIA11 (N=89) but are stationary with AcrIIA11 (N=93). P-value was determined by a Chi-Squared test (p = 3 × 10^−9^). (D) Lifetime of *Sa*Cas9 (N=30) and AcrIIA11:*Sa*Cas9 (N=49) on non-specific DNA. Half-lives are indicated for each curve. (E) Diffusion coefficients of *Sa*Cas9 with (N=33) and without (N=33) AcrIIA11. P-value was determined by a Mann-Whitney U-test (p = 9.6 × 10^−7^). Red diamonds indicate the mean diffusion coefficient.

To monitor the diffusion of wild-type *Sa*Cas9 RNPs, the sgRNA-complementary target was absent in this DNA substrate. The majority of *Sa*Cas9 RNPs (69%; N=61/89) scan the DNA via one-dimensional (1D) diffusion (**Figure 3B**). The remaining 31% of the molecules appeared stationary on DNA. *Sa*Cas9 RNPs are more diffusive than *Sp*Cas9, which only scans short stretches of DNA via 1D-diffusion (55). Half of the *Sa*Cas9 RNPs dissociated from the DNA within 61 ± 3 sec (N=30). Next, we pre-incubated unlabeled AcrIIA11 with *Sa*Cas9 and injected the AcrIIA11:*Sa*Cas9 complex to image its movement on DNA via the double-tethered DNA curtain assay. Only 25% of the AcrIIA11:*Sa*Cas9 molecules were diffusive (N=23/93), with the remaining 75% of the molecules appearing stationary (N=70/93) (**Figure 3C, Table S3**). AcrIIA11 also increased the lifetime of *Sa*Cas9 on non-specific DNA to 98 ± 4 sec (N=49) (**Figure 3D**). AcrIIA11 significantly decreased the *Sa*Cas9 diffusion coefficient. *Sa*Cas9 RNPs had a diffusion coefficient of 0.05 ± 0.01 µm^2^ s^-1^ (mean ± S.E.M.; N=33), whereas the AcrIIA11:*Sa*Cas9 complex had a mean diffusion coefficient of 0.006 ± 0.003 µm^2^ s^-1^ (N=33) (**Figure 3E, Table S3**). Since we cannot directly image AcrIIA11, these comparisons may underestimate the overall inhibition of diffusion, as any *Sa*Cas9 RNPs that are not complexed with AcrIIA11 are also considered in this analysis.

We confirmed these single-molecule results by assaying the relative DNA-binding affinities of nuclease dead *Sa*Cas9 (d*Sa*Cas9) with and without AcrIIA11 using electrophoretic mobility gel shift assays (EMSAs) (**Figure S4**). As expected, d*Sa*Cas9 weakly bound non-complementary DNA (11 ± 2% DNA bound) but had a higher affinity for the complementary DNA (52 ± 6% DNA bound). Increasing concentrations of AcrIIA11 increased d*Sa*Cas9 binding to the non-complementary DNA up to ∼3-fold (32 ± 3% bound DNA at 2 μM AcrIIA11). We previously showed AcrIIA11 is also a weak DNA-binding protein, but we did not detect any stable AcrIIA11 DNA binding at our working concentrations (**Figure S4A**) (22). Taken together, our single-molecule and ensemble biochemistry results indicate that AcrIIA11 slows the target search dynamics by retaining *Sa*Cas9 on off-target DNA sites.

To determine if AcrIIA11:*Sa*Cas9 blocked both target recognition and cleavage in the single-molecule assays, we loaded WT *Sa*Cas9 with a sgRNA targeting a single site in the middle of the DNA substrate (**Figure 4**). For these experiments, the digoxigenin DNA end was not immobilized, creating single-tethered DNA molecules in the presence of buffer flow, to visualize target binding and DNA cleavage. Injecting 2 nM WT *Sa*Cas9 RNPs resulted in 43% (N=38/89) of the molecules binding at the target site within 90 seconds of *Sa*Cas9 entering the flowcell (**Figure 4C-D, Table S3**). Most *Sa*Cas9s that did not localize to the target slid toward the free DNA end, consistent with buffer flow-induced biased diffusion (**Figure S5A**). Pre-incubating WT *Sa*Cas9 with AcrIIA11 resulted in a drastic decrease of target-bound *Sa*Cas9 molecules (12% target bound N=13/108 *Sa*Cas9s) (**Figure 4C-D, Table S3**). Of the 95 AcrIIA11:*Sa*Cas9 molecules not at the target, only 19% (N=18/95) slid to the DNA end. Instead, the majority of AcrIIA11:*Sa*Cas9 complexes bound non-specifically along the length of the DNA substrate. Similarly, d*Sa*Cas9 bound the target site (within our ∼500 bp spatial resolution) but was also trapped on non-specific DNA by AcrIIA11 (**Figure S5B-D**).

**Figure 4.**
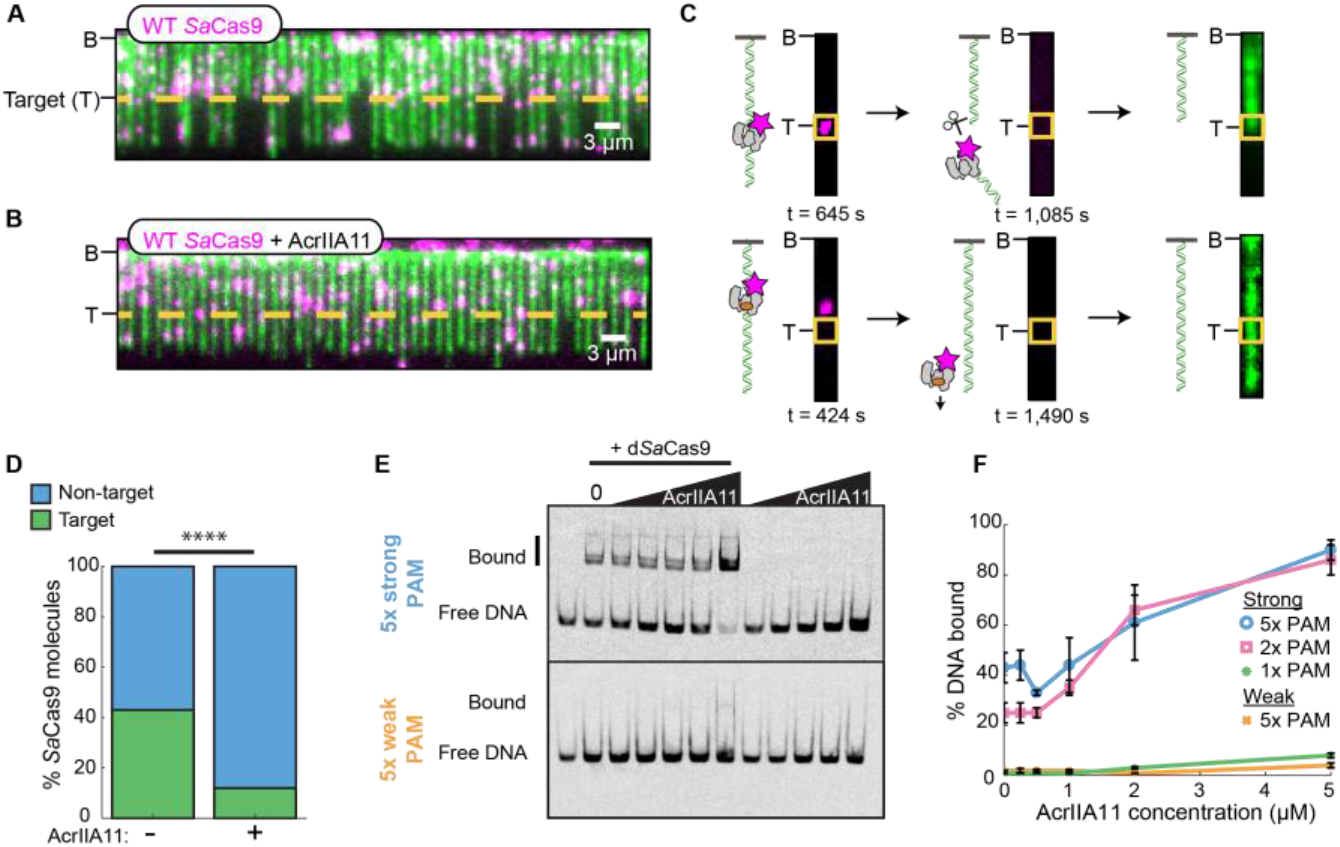
AcrIIA11 increases *Sa*Cas9 binding on PAM-rich non-specific DNA. (A) Image of single-tethered DNA molecules that are cleaved by WT *Sa*Cas9 RNPs. The target site (T) is indicated with a yellow dotted line. DNA is stained with a fluorescent intercalating dye YOYO-1 after incubation with *Sa*Cas9. (B) The DNA remains full-length after AcrIIA11:*Sa*Cas9 complexes are injected into the flowcell. (C) Schematic and images of *Sa*Cas9 binding (top, left), cleaving/ releasing the DNA (top, middle), and the cleaved DNA (top, right). Schematic and images of AcrIIA11: *Sa*Cas9 binding at an off-target site on the DNA (bottom, left), dissociating from the DNA (bottom, middle), and the remaining full-length DNA (bottom, right). The target site is labeled with a yellow box. (D) Percentage of WT *Sa*Cas9 at the target with (N=108) and without AcrIIA11 (N=89). P-value was determined by a Chi-Squared test (p = 1 × 10^−6^). (E) Representative EMSAs of the strong 5x PAM and weak 5x PAM DNA substrates with d*Sa*Cas9 and AcrIIA11. (F) Quantification of weak PAM, 1x, 2x, and 5x strong PAM EMSAs. Error bars: S.E.M. of three replicates.

Next, we analyzed the behavior of the AcrIIA11:*Sa*Cas9 complexes that appeared to bind the target site. *Sa*Cas9 is a multiple turnover enzyme that releases the PAM distal DNA after cleavage (56, 57). We designed the target DNA sequence so that *Sa*Cas9-catalyzed cleavage released both the enzyme and the DNA fragment from the tethered DNA end. Cleavage reactions could thus be determined by: (1) *Sa*Cas9 binding to the target site, (2) subsequent loss of the fluorescent *Sa*Cas9 signal, and (3) DNA cleavage at the target site, as confirmed by fluorescence imaging of the remaining tethered DNA (**Figure 4C**). These three criteria allowed us to rule out any *Sa*Cas9 molecules that bound the target site but dissociated without DNA cleavage. WT *Sa*Cas9s that stably bound the target cleaved and released 67% (N=20/30) of the DNA molecules (as imaged ∼15 minutes after RNP injection). As described above, we rarely observed AcrIIA11:*Sa*Cas9 complexes at the target DNA, but 40% (N=4/10) of those at the target site cleaved the DNA. We speculate that these four molecules may have not complexed with AcrIIA11 or happened to have reached the target site via random collisions. Nonetheless, we cannot rule out that AcrIIA11:*Sa*Cas9 complexes that reach the target site may retain cleavage activity.

Finally, we tested whether the AcrIIA11:*Sa*Cas9 complex prefers PAM-rich decoy sites. To test this, we incubated d*Sa*Cas9 with AcrIIA11 before adding sgRNA and non-target dsDNAs labeled on the 5’ end with a Cy5 fluorophore. We tested DNAs containing one, two, or five high-affinity PAMs (5’-NNGGGT). We also tested a DNA that contained five repeats of a low-affinity PAM (5’-NNTCTCN) (42). In contrast to *Sa*Cas9 RNPs, AcrIIA11:*Sa*Cas9 complexes demonstrated a strong preference for PAM sites (**Figure 4E-F and S5E-F**). Increasing the number of PAMs increased AcrIIA11:d*Sa*Cas9 binding, with 90% of the 5x strong PAM DNA bound at the highest AcrIIA11 concentration. There was a noticeable increase in AcrIIA11:d*Sa*Cas9 binding when we went from a single PAM to two PAMs, consistent with two *Sa*Cas9 RNPs in the AcrIIA11:*Sa*Cas9 complex. We conclude that the AcrIIA11:*Sa*Cas9 complex prefers PAM-rich decoy sites.

## Discussion

Here, we show that AcrIIA11 inhibits diverse Cas9 orthologs. We find AcrIIA11 inhibits *Sa*Cas9 cleavage by slowing facilitated diffusion and sequestering the AcrIIA11:*Sa*Cas9 complex at PAM-rich sites (**Figure 5**). Each *Sp*Cas9 searches *E. coli* for up to ∼6 hours to find its target, as it must query PAMs individually to check for target complementarity (58). By increasing Cas9 residence times at off-target sites, AcrIIA11 slows its facilitated diffusion and can delay CRISPR-based immunity sufficiently for the MGE to replicate. We also considered the possibility that AcrIIA11 can non-specifically coat DNA to protect it from *Sa*Cas9. However, we do not favor this inhibition mechanism for the following reasons: 1. AcrIIA11 directly interacts with Cas9; 2. AcrIIA11 inhibition is Cas9 specific (i.e., *Nme*Cas9 was not inhibited); 3. *Sa*Cas9 and *Sp*Cas9 could both bind DNA in the presence of AcrIIA11 (22). The mechanism outlined in **Figure 5** is similar to the one reported for AcrIF9, which also sequesters a type I-F Cascade complex on non-specific DNA (59, 60). Since both AcrIIA11 and AcrIF9 independently arrived at a similar “kinetic” inhibition mechanism for inactivating diverse Cas effectors, we propose that this is a successful strategy that nature has harnessed to inhibit additional CRISPR-Cas systems.

**Figure 5.**
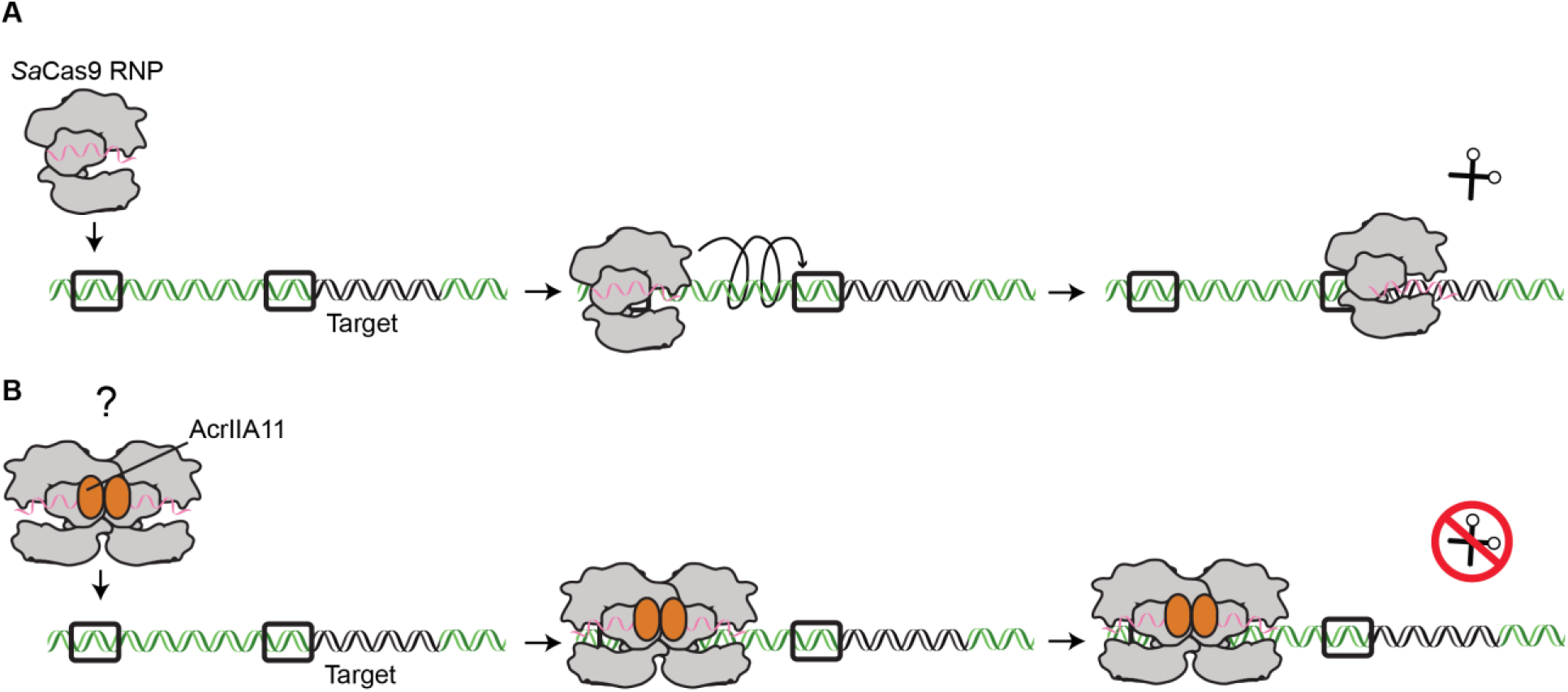
AcrIIA11 sequesters *Sa*Cas9 RNPs at non-specific DNA sites. (A) *Sa*Cas9 diffuses along the DNA in search of strong PAMs (black squares) that are adjacent to the target sequence. Once a possible target is recognized, *Sa*Cas9 forms an R-loop between the sgRNA and the target DNA, followed by DNA cleavage. (B) We speculate that AcrIIA11 dimerizes *Sa*Cas9 on DNA. The AcrIIA11:Cas9 complex is sequestered at off-target PAM-rich sites, preventing *Sa*Cas9 from recognizing and cleaving the target site.

Many Acrs dimerize or oligomerize their Cas effector complexes as part of the inhibition mechanism. For example, AcrIIC3 dimerizes *Nme*Cas9 and sterically blocks DNA binding (49, 61). However, dimerization of Cas12a by AcrVA4 is not required for inhibition (47). While the functional role of AcrVA4 dimerization of Cas12a is unknown, Wiegand et al. proposed that dimeric AcrVA4 may regulate its expression or that of another Acr (9). AcrIIA11 elutes as a dimer from a size exclusion column (22). We also note that AcrIIA11:d*Sa*Cas9 strongly prefers targets with two or more PAMs. Therefore, we favor a model where AcrIIA11 assembles a dimer of Cas9s and that this dimerization further increases non-specific DNA binding affinity. High-resolution structures of the AcrIIA11 dimer and the AcrIIA11:Cas9 complex will be required to further probe the role of the dimerization during inhibition.

Finally, we evaluated AcrIIA11 as a Cas9 inhibitor in mammalian cells. AcrIIA11 completely inhibited cleavage at three of four *Sa*Cas9-targeted loci. However, *Sa*Cas9 was only partially inhibited at the *RUNX1* site. We also observed incomplete inhibition of the *EMX1* locus when using *Sp*Cas9. Our biochemical mechanism hints at a possible reason for partial site-specific inhibition. We propose that these sites are more accessible to the nuclease and are thus cleaved before sufficient AcrIIA11 is expressed to complex with all available *Sa*Cas9 molecules. Recent work has shown Cas9 must bend the DNA to probe the sequence for target binding, so chromatin state and GC content may explain the differences in accessibility at these target sites (62). We note the overall indel percentage was highest at *RUNX1* compared to the other loci, suggesting that this site is more accessible to *Sa*Cas9. Alternatively, the AcrIIA11 and Cas9 may exist in a dynamic equilibrium where some of the complexes dissociate to transiently unleash the nuclease. Adjusting the timing of AcrIIA11 expression, the concentration of AcrIIA11 relative to Cas9, or the Cas9 ortholog used could potentially overcome this site-specific inhibition.

We find AcrIIA11 is a broad inhibitor of Cas9 orthologs (*Sp*Cas9, *Sa*Cas9, and *Fn*Cas9). Acrs that inhibit multiple Cas9 orthologs can be used as master inhibitors of Cas9 genome editing, making them versatile for many biotechnological applications. Other broad Cas9 inhibitors (AcrIIA5, AcrIIA16, AcrIIA17) either cleave the sgRNA or alter sgRNA expression levels, resulting in irreversible Cas9 inhibition (63–65). By not enzymatically altering Cas9, AcrIIA11 inhibition should be reversible. Further analysis of AcrIIA11 binding kinetics is required to determine the rate of reversibility of this inhibition, and mutations to the interface between Cas9 and AcrIIA11 could further fine-tune this inhibition and allow AcrIIA11 to act as an off- and on-switch for Cas9 cleavage. We conclude that AcrIIA11 is a broad inhibitor of Cas9s and inhibits via a reversible mechanism, making AcrIIA11 a promising master regulator of Cas9 genome editing.

## Materials and Methods

### Protein Cloning and Purification

*Sa*Cas9 (Addgene #101086) and AcrIIA11 were cloned into a pET19 expression vector containing an N-terminal 6xHis-TwinStrep-SUMO fusion (31). *Sa*Cas9 encoded an N-terminal 3xFLAG epitope for fluorescent labeling. Nuclease dead *Sa*Cas9 (d*Sa*Cas9) was created using primers KD197 and KD198 to mutate residues D10A and N580A (**Table S1**). AcrIIA11 used in pull-down assays was cloned into a pET19 expression vector with either a C-terminal TwinStrep (TS) or 6xHis epitope. SUMO protease was purified as previously described (32). For the AcrIIA11-TS pulldown of *Sa*Cas9 RNP, we cloned a sgRNA downstream of a T7 promoter into the original addgene #101086 plasmid (**Table S2**). For the TS-SUMO-*Sa*Cas9-sgRNA pulldown of AcrIIA11, the T7 promoter and sgRNA were cloned downstream of *Sa*Cas9 in the pET19 vector (**Table S2**). pMCSG7-Wt-NmeCas9 was a gift from Erik Sontheimer (Addgene plasmid #71474) (33). *Fn*Cas9 protein was purchased from Millipore Sigma (FNCAS9PROT-250UG).

*Sa*Cas9 was expressed in Rosetta (DE3) pLysS cells (VWR, 80509-788) and grown in 2 L LB supplemented with 100 µg/mL and 34µg/mL Carbenicillin and Chloramphenicol, respectively, at 37°C to an O.D._600_ ∼ 0.6. Cells were induced with 500 µM Isopropyl β-D-1-thiogalactopyranoside (IPTG) and grown overnight (∼16 hours) at 18°C. After induction, cells were pelleted via centrifugation and resuspended in Lysis Buffer (200 mM NaCl, 25 mM HEPES, pH 7.5, 2 mM DTT) with protease inhibitors and DNase before lysing with sonication. Cellular debris was pelleted by ultracentrifugation before placing lysate over a Strep-Tactin Superflow 50% suspension (IBA Life Sciences, 2-1206-025) gravity column equilibrated in Lysis Buffer. The column was washed with 100 mL of Lysis Buffer and eluted with 20 mL of Elution Buffer (200 mM NaCl, 25 mM HEPES, pH 7.5, 5 mM Desthiobiotin, 2 mM DTT). The eluate was concentrated with a 50 kDa Amicon Ultra-15 Centrifugal Filter (Millipore Sigma, UFC905096) and incubated with SUMO protease at 4°C overnight (∼16 hours). *Sa*Cas9 was isolated using a HiLoad 16/600 Superdex 200 pg column (Cytiva, 28989335) equilibrated in SEC Buffer (200 mM NaCl, 25 mM HEPES, pH 7.5, 5% glycerol, 2 mM DTT). Peak fractions were concentrated and frozen with liquid nitrogen before storing at -80°C.

AcrIIA11 was expressed in BL21 (DE3) RIL cells and grown in either LB or TB supplemented with 100 µg/mL and 34µg/mL Carbenicillin and Chloramphenicol, respectively, at 37°C to an O.D._600_ ∼ 0.6. Cells were induced with 200 µM IPTG for ∼16 hours at 18 °C. Cells were pelleted and resuspended in AcrIIA11 Lysis Buffer (500 mM NaCl, 25 mM HEPES, pH 7.5, 2 mM DTT) with protease inhibitors and DNase. Cells were lysed either via sonication or via the LM10 Microfluidizer before ultracentrifugation. Clarified lysate was placed on a Strep-Tactin Superflow 50% suspension (IBA Life Sciences, 2-1206-025) gravity column equilibrated in AcrIIA11 Lysis Buffer. The column was washed with 100 mL of AcrIIA11 Lysis Buffer and the protein was eluted with 20 mL of AcrIIA11 Elution Buffer (150 mM NaCl, 25 mM HEPES, pH 7.5, 5 mM Desthiobiotin, 2 mM DTT). The protein was incubated with SUMO protease at 4°C overnight before flowing over a Ni-NTA gravity column (ThermoFisher Scientific, 88222) to remove the cleaved 6xHis-TS-SUMO tag and SUMO Protease. The flow-through was concentrated to 1 mL using a 10 kDa Amicon Ultra-15 Centrifugal Filter (Millipore Sigma, UFC901096) before placing over a 5 mL Q column (Cytiva, 17515901) equilibrated in Q Buffer A (150mM NaCl, 25mM HEPES pH 7.5, 2mM DTT, 5% glycerol). The protein was eluted via a linear gradient with Q Buffer B (1M NaCl, 25mM HEPES, pH 7.5, 5% glycerol, 2mM DTT). Peak fractions were collected and concentrated to 1 mL and isolated on a HiLoad 16/600 Superdex 200 pg column (Cytiva, 28989335) equilibrated in AcrIIA11 SEC Buffer (200 mM NaCl, 25 mM HEPES, pH 7.5, 2mM DTT, 10% glycerol). Peak fractions were concentrated and frozen with liquid nitrogen before storing at -80°C.

*Nme*Cas9 was expressed in Rosetta 2(DE3) cells (Millipore Sigma, 71400-3) and grown in LB supplemented with 100 µg/mL Carbenicillin and 34µg/mL Chloramphenicol. Cultures were grown at 37°C to an O.D._600_ ∼ 0.6 and induced with 500 µM IPTG for ∼16 hours at 18°C. Cells were harvested by centrifugation at 6000xg for 15 minutes using a JLA 8.1000 rotor and resuspended in *Nme*Cas9 Lysis Buffer (500 mM NaCl, 50 mM Tris-HCL, pH 8, 10% glycerol, 1 mM TCEP) with protease inhibitors and DNase. Cells were lysed by sonication and pelleted using ultracentrifugation. Clarified lysate was placed on a 5 mL Ni-NTA gravity column and washed with 100 mL of *Nme*Cas9 Wash Buffer (500 mM NaCl, 50 mM Tris-HCL, pH 8, 10% glycerol, 1 mM TCEP, 25 mM Imidazole). *Nme*Cas9 was eluted with 15 mL of NmeCas9 Elution Buffer (500 mM NaCl, 50 mM Tris-HCL, pH 8, 10% glycerol, 1 mM TCEP, 200 mM Imidazole). The protein was dialyzed overnight (∼16 hours) at 4 °C with TEV protease into *Nme*Cas9 Dialysis Buffer (150 mM KCl, 20 mM HEPES, pH 7.5, 5% glycerol, 1 mM DTT). The sample was concentrated to 2 mL with a 30 kDa Amicon Ultra-15 Centrifugal Filter (Millipore Sigma, UFC903096) and placed on a 5 mL Heparin column (Cytiva, 17040703) equilibrated in *Nme*Cas9 Heparin A Buffer (150 mM KCl, 20 mM HEPES, pH 7.5, 5% glycerol, 1 mM DTT). The protein was eluted by a linear gradient with *Nme*Cas9 Heparin B Buffer (1 M KCl, 20 mM HEPES, pH 7.5, 5% glycerol, 1 mM DTT). The peak fractions were collected and spin concentrated to 1 mL before placing on the HiLoad 16/600 Superdex 200 pg column (Cytiva, 28989335) equilibrated in *Nme*Cas9 SEC Buffer (150 mM KCl, 20 mM HEPES, pH 7.5, 5% glycerol, 1 mM DTT). Peak fractions were collected, spin concentrated and frozen with liquid nitrogen before storing at -80°C.

### Human Cell Culture Genome Editing

The *Sa*Cas9 and CMV promoter-driven AcrIIA4 expression vectors were purchased from Addgene (Plasmid #85452 and #113038) (34, 35). Previously published sgRNAs for *CACNA1D, EMX1, FANCF*, and *RUNX1* (**Table S1**) were incorporated into the *Sa*Cas9 expression vector using Golden Gate cloning via Esp3I cut sites (36). The AcrIIA4 expression vector was modified to express AcrIIA11 with a C-terminal NLS and HA tags using the HiFi Assembly Kit (NEB).

HEK293T cells were cultured in DMEM (Thermo Fisher/Gibco) containing phenol red, 4 mM L-glutamine, 110 mg/L sodium pyruvate, 4.5 g/L D-glucose, and supplemented with 10% (v/v) FBS (Thermo Fisher/Gibco) and 100 U/mL penicillin + 100 μg/mL streptomycin (Thermo Fisher/Gibco). Cell lines were tested for mycoplasma contamination via the Mycoplasma Detection Kit (Southern Biotech). Transient transfections were performed with Lipofectamine 2000 (Life Technologies). Approximately 350,000 cells were seeded in each well of a 12-well plate 24 hr before transfection. Wells were transfected with either the Acr expression vector, the *Sa*Cas9/sgRNA expression vector, or both using 500 ng for each vector (3:1 Acr:*Sa*Cas9/sgRNA plasmid ratio) and either 1.5 µL or 3 µL of Lipofectamine (3 μL per μg of DNA).

HEK293T cells were collected and pelleted 72 hr post-transfection for genomic DNA extraction using the Wizard Genomic DNA Purification Kit (Promega). The target locus was PCR-amplified using AccuPrime Pfx high-fidelity DNA polymerase (Thermo Fisher) and the following PCR conditions: 95°C for 2 min, 35 cycles of 98°C for 15 s + 64°C for 30 s + 68°C for 2 min, and 68°C for 2 min. Reaction-specific primers are listed in (**Table S1**). Indel frequencies at the *Sa*Cas9 target site were assessed via a T7E1 assay with the EnGen Mutation Detection Kit (NEB), using the manufacturer’s recommendations. Reaction products were analyzed on a 1.3% SeaKem GTG agarose gel (Lonza) and imaged with the InGenuis3 (Syngene). For calculating indel percentages from gel images, bands from each lane were quantified with GelAnalyzer (version 2010a freeware). Peak areas for each band were measured and percentages of insertions and deletions [Indel(%)] were calculated using the formula: Indel(%)=100 × (1 – (1 – Fraction cleaved)*0.5), where Fraction cleaved = (Σ (Cleavage product bands))/(Σ (Cleavage product bands + PCR input band)).

### SaCas9 RNP:AcrIIA11 Pull-Down Assays

Rosetta (DE3) pLysS cells containing TS-SUMO-*Sa*Cas9-sgRNA or TS-SUMO-*Sa*Cas9 and BL21 (DE3) RIL cells containing AcrIIA11-6xHis were grown as described above. Cell pellets were lysed, separately, via sonication and pelleted via ultracentrifugation. The TS-SUMO-*Sa*Cas9-sgRNA or TS-SUMO-*Sa*Cas9 cell lysate was applied to a 5 mL Strep-Tactin Superflow 50% suspension (IBA Life Sciences, 2-1206-025) gravity column equilibrated in Lysis Buffer containing 200 mM NaCl, 25 mM HEPES, pH 7.5, 2 mM DTT supplemented with protease inhibitors and DNase. Then, cell lysate containing AcrIIA11-6xHis was applied to the same column, and the column was washed with 50 mL of Lysis Buffer. The complex was eluted with 20 mL of Elution Buffer (200 mM NaCl, 25 mM HEPES, pH 7.5, 5 mM Desthiobiotin, 2 mM DTT) and spin concentrated with a 50 kDa Amicon Ultra-15 Centrifugal Filter (Millipore Sigma, UFC905096) to ∼700µL. The complex was applied to a Superose 6 Increase 10/300 GL (Cytiva, 29091596) equilibrated in SEC Buffer (200 mM NaCl, 25 mM HEPES, pH 7.5, 5% glycerol, 2 mM DTT). Peak fractions were concentrated with a 50 kDa Amicon Ultra-15 Centrifugal Filter and frozen in liquid nitrogen before storing at -80°C. For the AcrIIA11-TS immobilization and 6xHis-*Sa*Cas9-sgRNA or 6xHis-*Sa*Cas9 pulldown, AcrIIA11-TS was immobilized on Strep-Tactin resin before *Sa*Cas9 RNP or apo*Sa*Cas9 was applied to the column. The rest of the purification is the same as above except this complex was purified on a HiLoad 16/600 Superdex 200 pg column (Cytiva, 28989335) instead of the Superose 6 Increase. For AcrIIA11-TS pulldown of 6xHis-*Sa*Cas9, 25 mM Tris-HCl, pH 8 was used in place of 25 mM HEPES, pH 7.5 in the Lysis, Elution, and SEC buffers.

### dSaCas9 and AcrIIA11 EMSAs with DNA containing PAMs

DNA oligos with a 5’ Cy5 fluorescent label contained either 5x weak PAMs (KD203 and KD204), 1x strong (KD247 and KD248), 2x strong (KD245 and KD246), or 5x strong PAMs (KD201 and KD202) (**Table S1**). Fluorescent oligos were incubated with 2x unlabeled oligos in Duplex Reaction Buffer (50 mM Tris-HCl, pH 8, 100 mM NaCl) at 90°C for 3 minutes before slowly bringing to room temperature. Proteins were diluted in Dilution Buffer (200 mM NaCl, 25 mM HEPES, pH 7.5, 5% glycerol, 2 mM DTT). 25 nM d*Sa*Cas9 was incubated with AcrIIA11 (250 nM, 500 nM, 1 µM, 2 µM, and 5 µM) in Cleavage Buffer (20 mM Tris-HCl, pH 8, 5% glycerol, 100 mM KCl, 5 mM MgCl_2_, 1 mM DTT) at room temperature for 10 minutes. 25 nM sgRNA and 5 nM DNA were added to the reaction and incubated for 10 minutes at 37°C. Samples were placed on ice and 10 mg Orange G + 30% glycerol was added. 6% Native PAGE gels containing 5 mM MgCl_2_ and 5% glycerol were pre-run in running buffer (0.5x TBE, 5 mM MgCl_2_, 5% glycerol) at 80V for 30 minutes. Wells were cleaned and samples were loaded before running the gels at 150V for 30 minutes at 4°C. Gels were imaged on the Typhoon FLA 9500 imager (Cytiva) and bands were quantified using GelAnalyzer 2010a.

### In vitro Cas9 ortholog inhibition assays

WT Cas9 (*Sa*Cas9, *Nme*Cas9, *Fn*Cas9) and AcrIIA11 were diluted in Dilution Buffer (200 mM NaCl, 25 mM HEPES, pH 7.5, 5% glycerol, 2 mM DTT). 50 nM of Cas9 was incubated with 5 µM AcrIIA11 and Cleavage Buffer (20 mM Tris-HCl, pH 8, 5% glycerol, 100 mM KCl, 5 mM MgCl_2_, 1 mM DTT) for 10 minutes at room temperature. 50 nM sgRNA and 4 nM linearized plasmid DNA was added, and the samples were placed at 37°C. For each time point, the samples were added to Quench Buffer (4.5 mg Orange G, 10% glycerol, 0.1 mM EDTA, pH 8, 0.02% SDS, ∼2mg/mL proteinase K (ThermoFisher Scientific, EO0491)) and incubated at 52°C for 30 minutes. Samples were run on a 1.25% agarose gel for 30 minutes at 120V before post-staining the gels with ethidium bromide. Gels were imaged using InGenuis3 (Syngene) and bands were quantified using GelAnalyzer 2010a.

### In vitro transcription of NmeCas9 and FnCas9 sgRNA

gBlocks containing either the *Nme*Cas9 or *Fn*Cas9 sgRNA sequence were ordered from IDT. gBlocks were amplified with Q5 polymerase and primers that annealed to the gBlock ends (KD142, KD143, KD144) (**Table S1**). PCR products were run on a 2% agarose gel post-stained with SYBR safe stain (APExBIO, A8743). PCR products were gel extracted using a QIAquick Gel Extraction Kit (Qiagen, 28704) and samples were eluted with RNase-free water. sgRNA was *in vitro* transcribed using HiScribe T7 High Yield RNA Synthesis Kit (NEB, E2040S). Samples were incubated at 37°C for 16 hours before purifying the sgRNA with Invitrogen TRIzol Reagent (Thermo Fisher Scientific, 15-596-018).

*Sa*Cas9 sgRNA was purchased from Synthego.

### dSaCas9 and AcrIIA11 EMSA with non-target DNA

A gBlock (SaCas9 SMART Target, **Table S1**) was amplified with primers IF365 and IF460 containing a 5’-ATTO647N (**Table S1**). d*Sa*Cas9 and AcrIIA11 were diluted in Dilution Buffer (200 mM NaCl, 25 mM HEPES, pH 7.5, 5% glycerol, 2 mM DTT). 50 nM d*Sa*Cas9 and different concentrations of AcrIIA11 (250 nM, 500 nM, 1 µM, 1.5 µM, and 2 µM) were incubated in Cleavage Buffer (20 mM Tris-HCl, pH 8, 5% glycerol, 100 mM KCl, 5 mM MgCl_2_, 1 mM DTT) for 10 minutes at room temperature. 5 nM DNA and 50 nM sgRNA were added and the reactions were incubated for 10 minutes at 37°C. Samples were placed on ice and 10 mg Orange G + 30% glycerol was added. 6% Native PAGE gels containing 5 mM MgCl_2_ and 5% glycerol were pre-run in running buffer (0.5x TBE, 5 mM MgCl_2_, 5% glycerol) for 30 minutes at 80V. Wells were cleaned and samples were loaded into the gel before running it at 150V for 1 hour at 4°C. Gels were imaged on the Typhoon FLA 9500 imager (Cytiva) and quantified using GelAnalyzer 2010a.

### sgRNA EMSA

The sgRNA used in this EMSA is listed in **Table S1** (SMART target sgRNA) and was labeled on the 5’ end with ^32^P. To avoid dissociation of the *Sa*Cas9-sgRNA complex during the binding experiments, *Sa*Cas9-sgRNA EMSAs were prepared with an excess of *Sa*Cas9 (490nM) incubated with a ^32^P-sgRNA (0.1nM) and cold sgRNA (490nM) mixture to pre-form the RNP. To form the RNP, *Sa*Cas9 and ^32^P-sgRNA substrates were incubated in charging buffer (10mM Tris-HCl, pH 8, 5mM MgCl_2_, 0.2mM DTT) at 25°C for 25 minutes. The RNP was prepared at a 100 nM effective concentration before adding AcrIIA11 (0.05µM, 0.1µM, 0.2µM, 0.8µM, 1.6µM, 3.2µM) and binding buffer (20mM Tris-HCl, pH 8, 5% glycerol, 1mM NaCl, 1mM DTT) for 30 minutes at 37°C. Samples were added to loading dye (1 mM Tris-HCl, pH 8, 0.1 mM EDTA, pH 8, 0.1 µg bromophenol blue, 0.1 µg xylene cyanol FF, 5% glycerol). A 10% Native PAGE (0.5X TBE) was pre-run at 80V for 15 minutes in 0.5x TBE running buffer. Samples were loaded into the gel and ran at 110 V for 45 minutes. Gels were dried at 80°C for 2 hours and exposed overnight. EMSAs were visualized by phosphorimaging using the Typhoon FLA 9500 imager (Cytiva).

### Single-molecule fluorescence microscopy and data analysis

All single-molecule imaging was performed using a Nikon Ti-E microscope in a prism-TIRF configuration equipped with a motorized stage (Prior ProScan II H117). Microfluidic flowcells were held by a custom-built stage heater to maintain experiments at 37°C. The flowcell was illuminated with a 488 nm (Coherent) laser through a quartz prism (Tower Optical Co.).

Microfluidic flowcells were prepared according to previously published protocols (37, 38). Double-tethered DNA curtains were prepared with 40 µL of liposome stock solution (97.7% DOPC, 2.0% DOPE-mPEG2k, and 0.3% DOPE-biotin; Avanti #850375P, #880130P, #870273P, respectively) in 960 µL Lipids Buffer (10 mM Tris-HCl, pH 8, 100 mM NaCl) incubated in the flowcell for 30 minutes. Then, 50 µg µL-1 of goat anti-rabbit polyclonal antibody (ICL Labs, #GGHL-15A) diluted in Lipids Buffer was incubated in the flowcell for 10 minutes. The flowcell was washed with BSA Buffer (40 mM Tris–HCl, pH 8, 2 mM MgCl_2_, 1 mM DTT, 0.2 mg mL-1 BSA) and 1 µg L-1 of digoxigenin monoclonal antibody (Life Technologies, #700772) diluted in BSA Buffer was injected and incubated for 10 minutes. Streptavidin (0.1 mg mL-1 diluted in BSA Buffer) was injected into the flowcell for another 10 minutes. Finally, ∼12.5 ng µL-1 of the biotin- and dig-labeled DNA substrate was injected into the flowcell. The anti-rabbit antibody and digoxigenin antibody steps were omitted to prepare single-tethered DNA curtains.

To fluorescently stain DNA with YOYO-1, ∼1 nM YOYO-1 (ThermoFisher, #Y3601), 1000 units of catalase (Millipore Sigma, #C100), 70 units of glucose oxidase (Millipore Sigma, # G2133), and 1% glucose (w/v) was injected into the flowcell at the end of the experiment.

For *Sa*Cas9 diffusion experiments, double-tethered curtains were assembled as described above. *Sa*Cas9 was diluted in dilution buffer (25 mM HEPES, pH 7.5, 200 mM NaCl, 5% glycerol) and incubated with 2x sgRNA (SMART target sgRNA, **Table S1**) in Cas9 charging buffer (50 mM Tris-HCl, pH 8, 10 mM MgCl_2_) for 5-10 minutes at room temperature (∼25°C) before adding Monoclonal Anti-FLAG BioM2 antibody (Millipore Sigma, #F9291) and Qdot 705 Streptavidin Conjugate (ThermoFisher, #Q10161MP) and placing the reaction on ice for ∼5 minutes. The reaction was diluted in imaging buffer (40 mM Tris-HCl, pH 8, 2 mM MgCl_2_, 1 mM DTT, 0.2 mg mL-1 BSA, 5µM Biotin) with 50 mM NaCl to a final concentration of 0.5 nM *Sa*Cas9 and 1 nM sgRNA and injected on the microscope. The experiment was conducted in imaging buffer with 50 mM NaCl. For AcrIIA11:*Sa*Cas9 diffusion experiments, *Sa*Cas9 and AcrIIA11 were diluted in dilution buffer and incubated in Cas9 charging buffer for 10 minutes at room temperature before adding 2x sgRNA to the reaction for 5 minutes at room temperature. Reactions were then incubated for ∼5 minutes with Monoclonal Anti-FLAG BioM2 antibody and Qdot 705 Streptavidin Conjugate on ice before diluting in imaging buffer with 50 mM NaCl to a final concentration of 1 nM *Sa*Cas9, 2 nM sgRNA, and 100 nM AcrIIA11. The concentration of *Sa*Cas9 on double-tethered curtains was lowered relative to AcrIIA11:*Sa*Cas9 to limit the number of diffusing *Sa*Cas9s per DNA and make tracking individual molecules easier.

For the d*Sa*Cas9 binding distribution, single-tethered DNA curtains were assembled and d*Sa*Cas9 was diluted and incubated with sgRNA (λ target-29.4 kb, **Table S1**) as described above. d*Sa*Cas9 was diluted in imaging buffer (40 mM Tris-HCl, pH 8, 2 mM MgCl_2_, 1 mM DTT, 0.2 mg mL-1 BSA, 5µM Biotin) with 100 mM NaCl to a final concentration of 1 nM d*Sa*Cas9 and 2 nM sgRNA. d*Sa*Cas9 RNP was injected onto the flowcell and incubated on the microscope for 5 minutes before imaging. The experiment was conducted in imaging buffer with 100 mM NaCl. For the AcrIIA11:d*Sa*Cas9 binding histogram, d*Sa*Cas9, AcrIIA11, and sgRNA were diluted and incubated as described above and diluted in imaging buffer with 100 mM NaCl to a final concentration of 1 nM d*Sa*Cas9, 2 nM sgRNA, and 100 nM AcrIIA11. AcrIIA11:d*Sa*Cas9 was injected into the flowcell and incubated on the microscope for 5 minutes before imaging.

To determine the localization and cleavage rate of WT *Sa*Cas9, single-tethered curtains were assembled. *Sa*Cas9 was diluted and incubated with sgRNA (λ target-29.4 kb, **Table S1**) as described above. *Sa*Cas9 RNP was diluted in imaging buffer (40 mM Tris-HCl, pH 8, 2 mM MgCl_2_, 1 mM DTT, 0.2 mg mL-1 BSA, 5µM Biotin) with 50 mM NaCl to a final concentration of 2 nM *Sa*Cas9 and 4 nM sgRNA and injected on the microscope. AcrIIA11:*Sa*Cas9 was diluted and incubated with sgRNA as described above. The complex was diluted in imaging buffer with 50 mM NaCl to a final concentration of 2 nM *Sa*Cas9, 4 nM sgRNA, and 200 nM AcrIIA11.

For the stationary vs diffusing analysis, a molecule was considered stationary if it stayed within a stationary 6-pixel window around the molecule. Molecules counted in the analysis had to bind DNA for ∼5 seconds or longer. The position along the DNA was determined by measuring the distance of *Sa*Cas9 relative to the barrier. The position of WT *Sa*Cas9 with and without AcrIIA11 was determined 90 seconds after the molecules entered the flowcell. To determine if a DNA molecule was cleaved by *Sa*Cas9, *Sa*Cas9 had to bind the target and release the DNA (loss of *Sa*Cas9 signal). The DNA was stained at the end of the experiment with YOYO-1 to confirm it was cleaved. *Sa*Cas9 molecules that bound the target and did not cleave the DNA had to remain on the DNA for the rest of the movie without cleaving it to be counted in the cleavage analysis. Molecules that bound the target and dissociated during the movie without cleaving it were not counted. Binding lifetimes were fit to a single exponential decay using a custom MATLAB script (Mathworks R2017a).

For the diffusion coefficient analysis, particle trajectories were tracked using a custom ImageJ script. MSD was calculated for the first 10 time intervals of each particle and fit to a line to obtain the diffusion coefficient, as previously described (39). The mean diffusion coefficient was obtained from >30 molecules and the standard error of the mean (S.E.M.) was determined.

## End Matter

### Author Contributions and Notes

K.E.D., K.J.F., and I.J.F. conceived the study. K.E.D. and C.T. designed, performed, and analyzed experiments. K.E.D. generated key materials. W.K. conducted biochemical assays. K.J.F. established protocols for purification and some *in vitro* AcrIIA11 assays. K.J. designed, performed, and analyzed human cell line work.

The authors declare no conflict of interest.

## Funding

This work is supported by the Welch Foundation (F-1808 to I.J.F.) and the NIH (R01GM124141 to I.J.F., R01GM104896 to W.K., and F31GM125201 to K.E.D.). K.J.F. was supported by a Helen Hay Whitney Foundation postdoctoral fellowship, the Endowed Scholars Program at the University of Texas Southwestern Medical Center, and by NIH grant 1DP2-AI154402.

## Data Availability

Reprints and permissions information is available online. Correspondence and requests for materials should be addressed to K.E.D. and I.J.F.

## Notes

### Competing Interest Statement

The authors have declared no competing interest.

## References

1. 1.Jinek, M., Chylinski, K., Fonfara, I., Hauer, M., Doudna, J.A. and Charpentier, E. (2012) A Programmable Dual-RNA–Guided DNA Endonuclease in Adaptive Bacterial Immunity. Science, 337, 816–821.

2. Gasiunas, G., Barrangou, R., Horvath, P. and Siksnys, V. (2012) Cas9– crRNA ribonucleoprotein complex mediates specific DNA cleavage for adaptive immunity in bacteria. Proc. Natl. Acad. Sci., 109, E2579–E2586.

3. Singh, D., Sternberg, S.H., Fei, J., Doudna, J.A. and Ha, T. (2016) Real-time observation of DNA recognition and rejection by the RNA-guided endonuclease Cas9. Nat. Commun., 7, 12778.

4. Sternberg, S.H., Redding, S., Jinek, M., Greene, E.C. and Doudna, J.A. (2014) DNA interrogation by the CRISPR RNA-guided endonuclease Cas9. Nature, 507, 62–67.

5. Szczelkun, M.D., Tikhomirova, M.S., Sinkunas, T., Gasiunas, G., Karvelis, T., Pschera, P., Siksnys, V. and Seidel, R. (2014) Direct observation of R-loop formation by single RNA-guided Cas9 and Cascade effector complexes. Proc. Natl. Acad. Sci. U. S. A., 111, 9798–9803.

6. Jiang, F. and Doudna, J.A. (2017) CRISPR–Cas9 Structures and Mechanisms. Annu. Rev. Biophys., 46, 505–529.

7. Swartjes, T., Staals, R.H.J. and van der Oost, J. (2020) Editor’s cut: DNA cleavage by CRISPR RNA-guided nucleases Cas9 and Cas12a. Biochem. Soc. Trans., 48, 207–219.

8. Hwang, S. and Maxwell, K.L. (2019) Meet the Anti-CRISPRs: Widespread Protein Inhibitors of CRISPR-Cas Systems. CRISPR J., 2, 23–30.

9. Wiegand, T., Karambelkar, S., Bondy-Denomy, J. and Wiedenheft, B. (2020) Structures and Strategies of Anti-CRISPR-Mediated Immune Suppression. Annu. Rev. Microbiol., 74, 21–37.

10. Davidson, A.R., Lu, W.-T., Stanley, S.Y., Wang, J., Mejdani, M., Trost, C.N., Hicks, B.T., Lee, J. and Sontheimer, E.J. (2020) Anti-CRISPRs: Protein Inhibitors of CRISPR-Cas Systems. Annu. Rev. Biochem., 89, 309–332.

11. Bondy-Denomy, J., Pawluk, A., Maxwell, K.L. and Davidson, A.R. (2013) Bacteriophage genes that inactivate the CRISPR/Cas bacterial immune system. Nature, 493, 429–432.

12. Chowdhury, S., Carter, J., Rollins, M.F., Golden, S.M., Jackson, R.N., Hoffmann, C., Nosaka, L., Bondy-Denomy, J., Maxwell, K.L., Davidson, A.R., et al. (2017) Structure Reveals Mechanisms of Viral Suppressors that Intercept a CRISPR RNA-Guided Surveillance Complex. Cell, 169, 47-57.e11.

13. Bondy-Denomy, J., Garcia, B., Strum, S., Du, M., Rollins, M.F., Hidalgo-Reyes, Y., Wiedenheft, B., Maxwell, K.L. and Davidson, A.R. (2015) Multiple mechanisms for CRISPR–Cas inhibition by anti-CRISPR proteins. Nature, 526, 136–139.

14. Guo, T.W., Bartesaghi, A., Yang, H., Falconieri, V., Rao, P., Merk, A., Eng, E.T., Raczkowski, A.M., Fox, T., Earl, L.A., et al. (2017) Cryo-EM Structures Reveal Mechanism and Inhibition of DNA Targeting by a CRISPR-Cas Surveillance Complex. Cell, 171, 414-426.e12.

15. Wang, X., Yao, D., Xu, J.-G., Li, A.-R., Xu, J., Fu, P., Zhou, Y. and Zhu, Y. (2016) Structural basis of Cas3 inhibition by the bacteriophage protein AcrF3. Nat. Struct. Mol. Biol., 23, 868–870.

16. Pawluk, A., Staals, R.H.J., Taylor, C., Watson, B.N.J., Saha, S., Fineran, P.C., Maxwell, K.L. and Davidson, A.R. (2016) Inactivation of CRISPR-Cas systems by anti-CRISPR proteins in diverse bacterial species. Nat. Microbiol., 1, 1–6.

17. Pawluk, A., Amrani, N., Zhang, Y., Garcia, B., Hidalgo-Reyes, Y., Lee, J., Edraki, A., Shah, M., Sontheimer, E.J., Maxwell, K.L., et al. (2016) Naturally Occurring Off-Switches for CRISPR-Cas9. Cell, 167, 1829-1838.e9.

18. Rauch, B.J., Silvis, M.R., Hultquist, J.F., Waters, C.S., McGregor, M.J., Krogan, N.J. and Bondy-Denomy, J. (2017) Inhibition of CRISPR-Cas9 with Bacteriophage Proteins. Cell, 168, 150-158.e10.

19. Hynes, A.P., Rousseau, G.M., Lemay, M.-L., Horvath, P., Romero, D.A., Fremaux, C. and Moineau, S. (2017) An anti-CRISPR from a virulent streptococcal phage inhibits Streptococcus pyogenes Cas9. Nat. Microbiol., 2, 1374–1380.

20. Hynes, A.P., Rousseau, G.M., Agudelo, D., Goulet, A., Amigues, B., Loehr, J., Romero, D.A., Fremaux, C., Horvath, P., Doyon, Y., et al. (2018) Widespread anti-CRISPR proteins in virulent bacteriophages inhibit a range of Cas9 proteins. Nat. Commun., 9, 2919.

21. Watters, K.E., Fellmann, C., Bai, H.B., Ren, S.M. and Doudna, J.A. (2018) Systematic discovery of natural CRISPR-Cas12a inhibitors. Science, 362, 236–239.

22. Forsberg, K.J., Bhatt, I.V., Schmidtke, D.T., Javanmardi, K., Dillard, K.E., Stoddard, B.L., Finkelstein, I.J., Kaiser, B.K. and Malik, H.S. (2019) Functional metagenomics-guided discovery of potent Cas9 inhibitors in the human microbiome. eLife, 8, e46540.

23. Uribe, R.V., van der Helm, E., Misiakou, M.-A., Lee, S.-W., Kol, S. and Sommer, M.O.A. (2019) Discovery and Characterization of Cas9 Inhibitors Disseminated across Seven Bacterial Phyla. Cell Host Microbe, 25, 233-241.e5.

24. Shin, J., Jiang, F., Liu, J.-J., Bray, N.L., Rauch, B.J., Baik, S.H., Nogales, E., Bondy-Denomy, J., Corn, J.E. and Doudna, J.A. (2017) Disabling Cas9 by an anti-CRISPR DNA mimic. Sci. Adv., 3, e1701620.

25. Li, C., Psatha, N., Gil, S., Wang, H., Papayannopoulou, T. and Lieber, A. (2018) HDAd5/35++ Adenovirus Vector Expressing Anti-CRISPR Peptides Decreases CRISPR/Cas9 Toxicity in Human Hematopoietic Stem Cells. Mol. Ther. Methods Clin. Dev., 9, 390–401.

26. Lee, J., Mir, A., Edraki, A., Garcia, B., Amrani, N., Lou, H.E., Gainetdinov, I., Pawluk, A., Ibraheim, R., Gao, X.D., et al. (2018) Potent Cas9 Inhibition in Bacterial and Human Cells by AcrIIC4 and AcrIIC5 Anti-CRISPR Proteins. mBio, 9.

27. Lee, J., Mou, H., Ibraheim, R., Liang, S.-Q., Liu, P., Xue, W. and Sontheimer, E.J. (2019) Tissue-restricted genome editing in vivo specified by microRNA-repressible anti-CRISPR proteins. RNA N. Y. N, 25, 1421–1431.

28. Hoffmann, M.D., Aschenbrenner, S., Grosse, S., Rapti, K., Domenger, C., Fakhiri, J., Mastel, M., Börner, K., Eils, R., Grimm, D., et al. (2019) Cell-specific CRISPR-Cas9 activation by microRNA-dependent expression of anti-CRISPR proteins. Nucleic Acids Res., 47, e75.

29. Matsumoto, D., Tamamura, H. and Nomura, W. (2020) A cell cycle-dependent CRISPR-Cas9 activation system based on an anti-CRISPR protein shows improved genome editing accuracy. Commun. Biol., 3, 1–10.

30. Nakamura, M., Srinivasan, P., Chavez, M., Carter, M.A., Dominguez, A.A., La Russa, M., Lau, M.B., Abbott, T.R., Xu, X., Zhao, D., et al. (2019) Anti-CRISPR-mediated control of gene editing and synthetic circuits in eukaryotic cells. Nat. Commun., 10, 194.

31. Soares Medeiros, L.C., South, L., Peng, D., Bustamante, J.M., Wang, W., Bunkofske, M., Perumal, N., Sanchez-Valdez, F. and Tarleton, R.L. (2017) Rapid, Selection-Free, High-Efficiency Genome Editing in Protozoan Parasites Using CRISPR-Cas9 Ribonucleoproteins. mBio, 8, e01788–17.

32. Malakhov, M.P., Mattern, M.R., Malakhova, O.A., Drinker, M., Weeks, S.D. and Butt, T.R. (2004) SUMO fusions and SUMO-specific protease for efficient expression and purification of proteins. J. Struct. Funct. Genomics, 5, 75–86.

33. Zhang, Y., Rajan, R., Seifert, H.S., Mondragón, A. and Sontheimer, E.J. (2015) DNase H Activity of Neisseria meningitidis Cas9. Mol. Cell, 60, 242–255.

34. Bubeck, F., Hoffmann, M.D., Harteveld, Z., Aschenbrenner, S., Bietz, A., Waldhauer, M.C., Börner, K., Fakhiri, J., Schmelas, C., Dietz, L., et al. (2018) Engineered anti-CRISPR proteins for optogenetic control of CRISPR–Cas9. Nat. Methods, 15, 924–927.

35. Singer, M., Wang, C., Cong, L., Marjanovic, N.D., Kowalczyk, M.S., Zhang, H., Nyman, J., Sakuishi, K., Kurtulus, S., Gennert, D., et al. (2016) A Distinct Gene Module for Dysfunction Uncoupled from Activation in Tumor-Infiltrating T Cells. Cell, 166, 1500-1511.e9.

36. Kleinstiver, B.P., Prew, M.S., Tsai, S.Q., Nguyen, N.T., Topkar, V.V., Zheng, Z. and Joung, J.K. (2015) Broadening the targeting range of Staphylococcus aureus CRISPR-Cas9 by modifying PAM recognition. Nat. Biotechnol., 33, 1293–1298.

37. Gallardo, I.F., Pasupathy, P., Brown, M., Manhart, C.M., Neikirk, D.P., Alani, E. and Finkelstein, I.J. (2015) High-Throughput Universal DNA Curtain Arrays for Single-Molecule Fluorescence Imaging. Langmuir, 31, 10310–10317.

38. Soniat, M.M., Myler, L.R., Schaub, J.M., Kim, Y., Gallardo, I.F. and Finkelstein, I.J. (2017) Next-Generation DNA Curtains for Single-Molecule Studies of Homologous Recombination. Methods Enzymol., 592, 259–281.

39. Brown, M.W., Kim, Y., Williams, G.M., Huck, J.D., Surtees, J.A. and Finkelstein, I.J. (2016) Dynamic DNA binding licenses a repair factor to bypass roadblocks in search of DNA lesions. Nat. Commun., 7, 10607.

40. Acharya, S., Mishra, A., Paul, D., Ansari, A.H., Azhar, M., Kumar, M., Rauthan, R., Sharma, N., Aich, M., Sinha, D., et al. (2019) Francisella novicida Cas9 interrogates genomic DNA with very high specificity and can be used for mammalian genome editing. Proc. Natl. Acad. Sci., 116, 20959–20968.

41. Nishimasu, H., Cong, L., Yan, W.X., Ran, F.A., Zetsche, B., Li, Y., Kurabayashi, A., Ishitani, R., Zhang, F. and Nureki, O. (2015) Crystal Structure of Staphylococcus aureus Cas9. Cell, 162, 1113–1126.

42. Ran, F.A., Cong, L., Yan, W.X., Scott, D.A., Gootenberg, J.S., Kriz, A.J., Zetsche, B., Shalem, O., Wu, X., Makarova, K.S., et al. (2015) In vivo genome editing using Staphylococcus aureus Cas9. Nature, 520, 186–191.

43. Song, G., Zhang, F., Zhang, X., Gao, X., Zhu, X., Fan, D. and Tian, Y. (2019) AcrIIA5 Inhibits a Broad Range of Cas9 Orthologs by Preventing DNA Target Cleavage. Cell Rep., 29, 2579-2589.e4.

44. Dong, D., Guo, M., Wang, S., Zhu, Y., Wang, S., Xiong, Z., Yang, J., Xu, Z. and Huang, Z. (2017) Structural basis of CRISPR–SpyCas9 inhibition by an anti-CRISPR protein. Nature, 546, 436–439.

45. Fuchsbauer, O., Swuec, P., Zimberger, C., Amigues, B., Levesque, S., Agudelo, D., Duringer, A., Chaves-Sanjuan, A., Spinelli, S., Rousseau, G.M., et al. (2019) Cas9 Allosteric Inhibition by the Anti-CRISPR Protein AcrIIA6. Mol. Cell, 76, 922-937.e7.

46. Ka, D., An, S.Y., Suh, J.-Y. and Bae, E. (2018) Crystal structure of an anti-CRISPR protein, AcrIIA1. Nucleic Acids Res., 46, 485–492.

47. Knott, G.J., Cress, B.F., Liu, J.-J., Thornton, B.W., Lew, R.J., Al-Shayeb, B., Rosenberg, D.J., Hammel, M., Adler, B.A., Lobba, M.J., et al. Structural basis for AcrVA4 inhibition of specific CRISPR-Cas12a. eLife, 8, e49110.

48. Knott, G.J., Thornton, B.W., Lobba, M.J., Liu, J.-J., Al-Shayeb, B., Watters, K.E. and Doudna, J.A. (2019) Broad-spectrum enzymatic inhibition of CRISPR-Cas12a. Nat. Struct. Mol. Biol., 26, 315–321.

49. Zhu, Y., Gao, A., Zhan, Q., Wang, Y., Feng, H., Liu, S., Gao, G., Serganov, A. and Gao, P. (2019) Diverse Mechanisms of CRISPR-Cas9 Inhibition by Type IIC Anti-CRISPR Proteins. Mol. Cell, 74, 296-309.e7.

50. Peng, R., Xu, Y., Zhu, T., Li, N., Qi, J., Chai, Y., Wu, M., Zhang, X., Shi, Y., Wang, P., et al. (2017) Alternate binding modes of anti-CRISPR viral suppressors AcrF1/2 to Csy surveillance complex revealed by cryo-EM structures. Cell Res., 27, 853–864.

51. Liu, L., Yin, M., Wang, M. and Wang, Y. (2019) Phage AcrIIA2 DNA Mimicry: Structural Basis of the CRISPR and Anti-CRISPR Arms Race. Mol. Cell, 73, 611-620.e3.

52. Peng, R., Li, Z., Xu, Y., He, S., Peng, Q., Wu, L., Wu, Y., Qi, J., Wang, P., Shi, Y., et al. (2019) Structural insight into multistage inhibition of CRISPR-Cas12a by AcrVA4. Proc. Natl. Acad. Sci., 116, 18928–18936.

53. Thavalingam, A., Cheng, Z., Garcia, B., Huang, X., Shah, M., Sun, W., Wang, M., Harrington, L., Hwang, S., Hidalgo-Reyes, Y., et al. (2019) Inhibition of CRISPR-Cas9 ribonucleoprotein complex assembly by anti-CRISPR AcrIIC2. Nat. Commun., 10, 2806.

54. Zhang, H., Li, Z., Daczkowski, C.M., Gabel, C., Mesecar, A.D. and Chang, L. (2019) Structural Basis for the Inhibition of CRISPR-Cas12a by Anti-CRISPR Proteins. Cell Host Microbe, 25, 815-826.e4.

55. Globyte, V., Lee, S.H., Bae, T., Kim, J.-S. and Joo, C. (2019) CRISPR/Cas9 searches for a protospacer adjacent motif by lateral diffusion. EMBO J., 38, e99466.

56. Zhang, S., Zhang, Q., Hou, X.-M., Guo, L., Wang, F., Bi, L., Zhang, X., Li, H.-H., Wen, F., Xi, X.-G., et al. (2020) Dynamics of Staphylococcus aureus Cas9 in DNA target Association and Dissociation. EMBO Rep., 21, e50184.

57. Yourik, P., Fuchs, R.T., Mabuchi, M., Curcuru, J.L. and Robb, G.B. (2019) Staphylococcus aureus Cas9 is a multiple-turnover enzyme. RNA, 25, 35–44.

58. Jones, D.L., Leroy, P., Unoson, C., Fange, D., Ćurić, V., Lawson, M.J. and Elf, J. (2017) Kinetics of dCas9 target search in Escherichia coli. Science, 357, 1420–1424.

59. Hirschi, M., Lu, W.-T., Santiago-Frangos, A., Wilkinson, R., Golden, S.M., Davidson, A.R., Lander, G.C. and Wiedenheft, B. (2020) AcrIF9 tethers non-sequence specific dsDNA to the CRISPR RNA-guided surveillance complex. Nat. Commun., 11, 2730.

60. Lu, W.-T., Trost, C.N., Müller-Esparza, H., Randau, L. and Davidson, A.R. (2021) Anti-CRISPR AcrIF9 functions by inducing the CRISPR–Cas complex to bind DNA non-specifically. Nucleic Acids Res., 49, 3381–3393.

61. Harrington, L.B., Doxzen, K.W., Ma, E., Liu, J.-J., Knott, G.J., Edraki, A., Garcia, B., Amrani, N., Chen, J.S., Cofsky, J.C., et al. (2017) A Broad-Spectrum Inhibitor of CRISPR-Cas9. Cell, 170, 1224-1233.e15.

62. Cofsky, J.C., Soczek, K.M., Knott, G.J., Nogales, E. and Doudna, J.A. (2021) CRISPR-Cas9 bends and twists DNA to read its sequence.

63. An, S.Y., Ka, D., Kim, I., Kim, E.-H., Kim, N.-K., Bae, E. and Suh, J.-Y. (2020) Intrinsic disorder is essential for Cas9 inhibition of anti-CRISPR AcrIIA5. Nucleic Acids Res., 48, 7584–7594.

64. Garcia, B., Lee, J., Edraki, A., Hidalgo-Reyes, Y., Erwood, S., Mir, A., Trost, C.N., Seroussi, U., Stanley, S.Y., Cohn, R.D., et al. (2019) Anti-CRISPR AcrIIA5 Potently Inhibits All Cas9 Homologs Used for Genome Editing. Cell Rep., 29, 1739-1746.e5.

65. Mahendra, C., Christie, K.A., Osuna, B.A., Pinilla-Redondo, R., Kleinstiver, B.P. and Bondy-Denomy, J. (2020) Broad-spectrum anti-CRISPR proteins facilitate horizontal gene transfer. Nat. Microbiol., 5, 620–629.

