## Supplemental data for "Mechanism of broad-spectrum Cas9 inhibition by AcrIIA11"

### Supplemental Figures

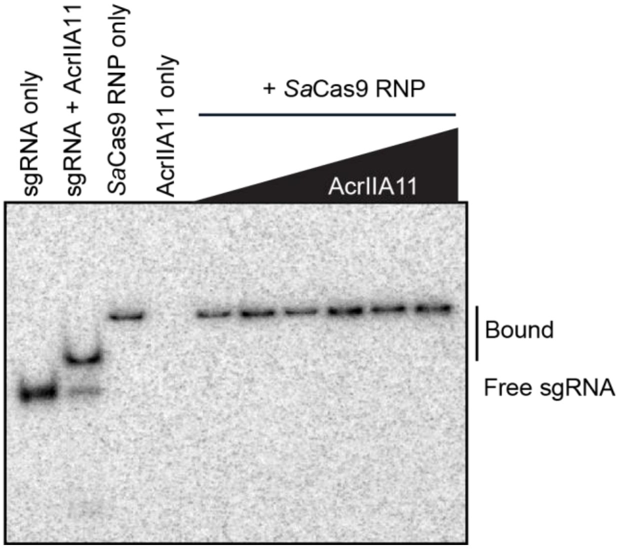


**Figure S1. AcrIIA11 binds but does not degrade sgRNA.**

Native PAGE gel of $^{32}$P-labeled sgRNA bound by *Sa*Cas9 and incubated with various concentrations of AcrIIA11 for 30 minutes at 37˚C. AcrIIA11 weakly binds sgRNA in the absence of *Sa*Cas9.

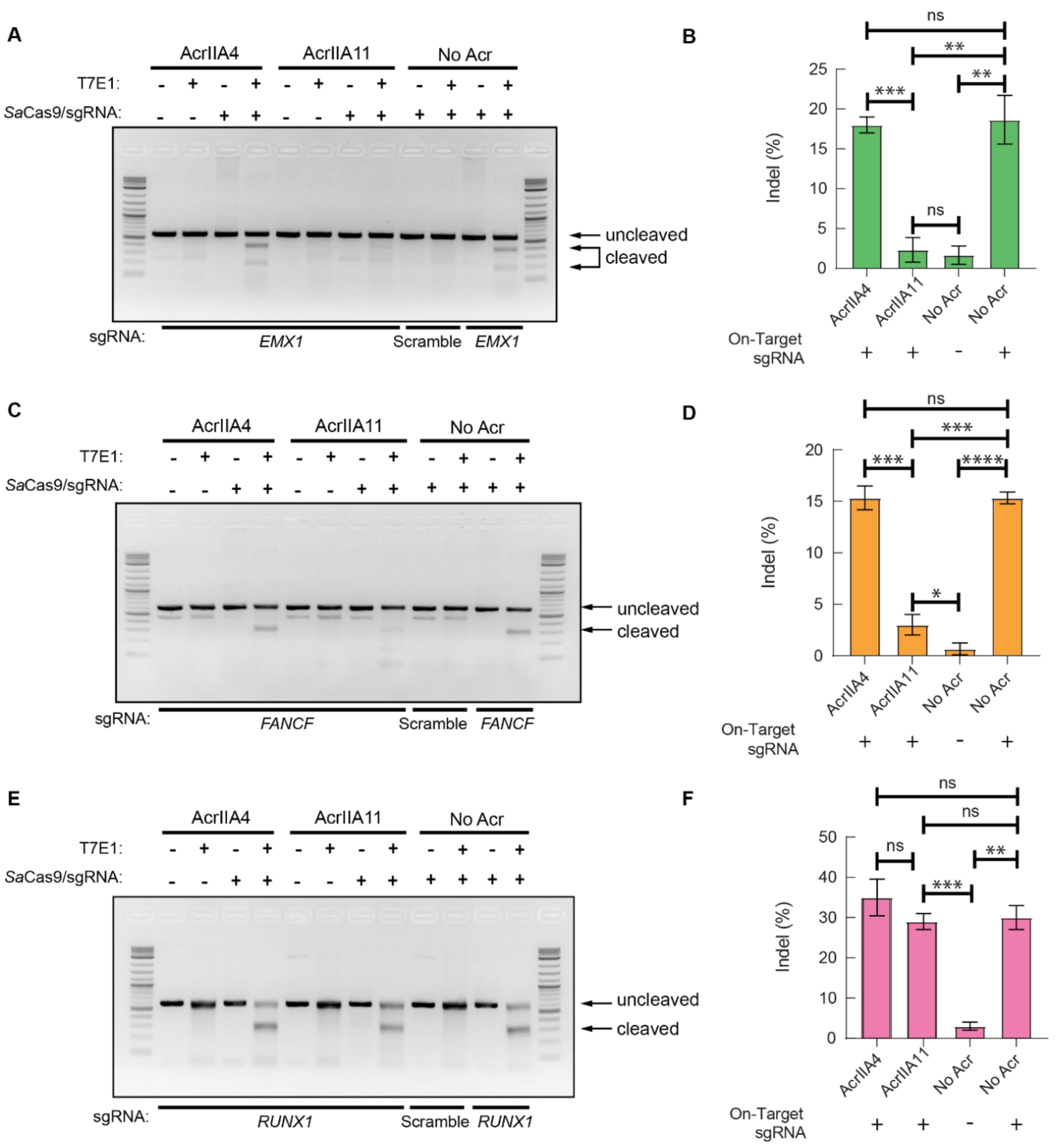


**Figure S2. AcrIIA11 inhibits *Sa*Cas9 cleavage in human cells.**

Representative agarose gels showing *Sa*Cas9 genome editing and the quantification of three replicates at the (A, B) *EMX1*, (C, D) *FANCF*, and (E, F) *RUNX1* sites with or without AcrIIA11. Error bars are standard deviation of three replicates. P-values (not significant [ns], $p > 0.05$; $*p < 0.05$; $**p < 0.01$; $***p < 0.001$; $****p < 0.0001$) were determined using a Student's t-test.

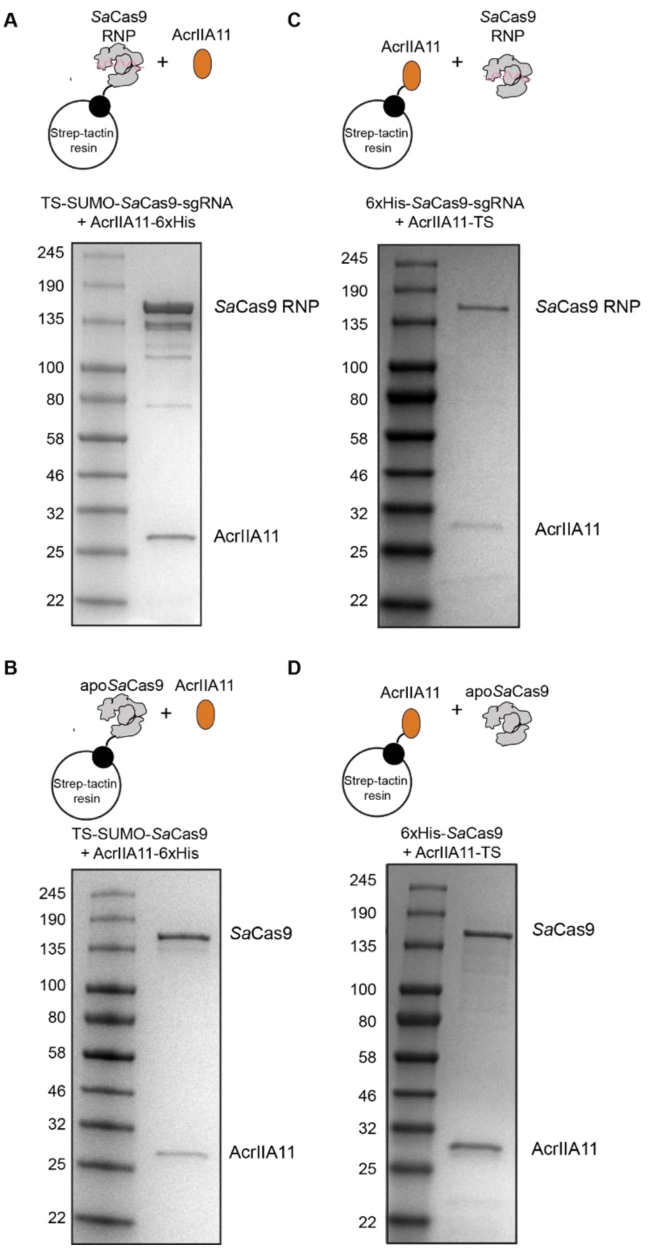


**Figure S3. *Sa*Cas9 physically interacts with AcrIIA11.**

(A, B) Schematic and Coomassie-stained SDS-PAGE gel of AcrIIA11 pulldown. *Sa*Cas9 (A) RNP or apo*Sa*Cas9 (B) were immobilized on Strep-Tactin resin. All AcrIIA11:*Sa*Cas9 complexes were then purified on a size exclusion column (SEC). (C,D) Strep-Tactin-immobilized AcrIIA11 can also pulldown *Sa*Cas9 RNP (C) and apo*Sa*Cas9 (D).

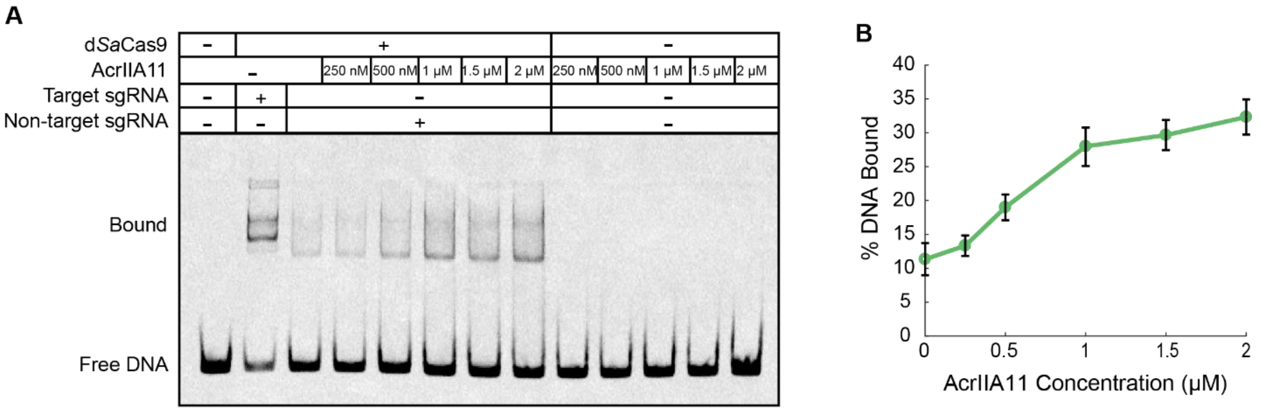


**Figure S4. AcrIIA11 induces non-specific binding of *Sa*Cas9 on DNA.**

(A) EMSA of d*Sa*Cas9 non-specific binding at increasing concentrations of AcrIIA11. AcrIIA11 alone does not stably bind the DNA at these concentrations. (B) Quantification of three EMSA replicates. Error bars are S.E.M.

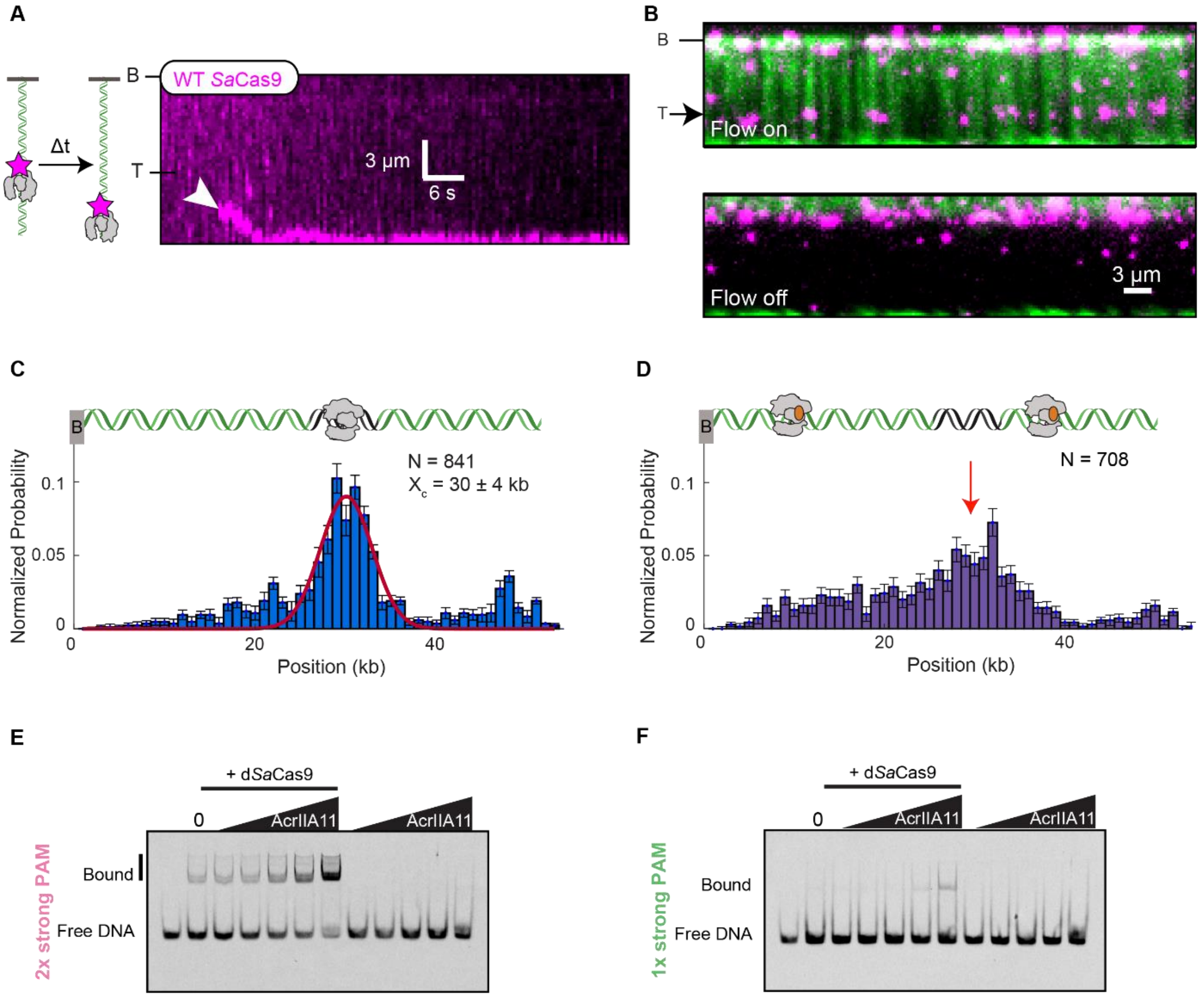


**Figure S5. AcrIIA11 sequesters *Sa*Cas9 at off-target PAMs sites.**

(A) Schematic and kymograph showing WT *Sa*Cas9 sliding down to the DNA end. The white arrow indicates *Sa*Cas9 binding. (B) Images of d*Sa*Cas9 at the target sequence. Top: buffer flow is on. Bottom: buffer flow is off. DNA retracts to the barrier. (C) Binding histogram of d*Sa*Cas9 binding to the target site at 29.4 kb. Fit to a single Gaussian (center and SD are indicated). The second peak indicates molecules that slide to the free DNA end, as shown in panel (A). (D) Binding histogram of AcrIIA11:d*Sa*Cas9. The red arrow indicates the target site. (E, F) Representative EMSAs of AcrIIA11:d*Sa*Cas9 binding to the (E) 2x strong and (F) 1x strong PAM DNA.

**Table S1. Oligonucleotides, sgRNA, and gBlocks used in this study.**

| Oligonucleotide | Sequence |
|---|---|
| KD197 | AACTACATTCTGGGGCTGGCCATCGGGATTACAAGCGTG |
| KD198 | TGCTGGTCAAGCAGGAAGAGGCATCTAAAAAGGGCAATAGGAC |
| KD201 | /5Cy5/tttgggtattgggtattgggttttgggtttgggttatttt |
| KD202 | aaaataacccaaacccaaaacccaatacccaatacccaaa |
| KD203 | /5Cy5/tttTCTCattTCTCattTCTCtttTCTCttTCTCtatttt |
| KD204 | aaaataGAGAaaGAGAaaaGAGAaatGAGAaatGAGAaaa |
| KD245 | /5Cy5/ tttgggtattTCTCattTCTCtttTCTCttgggttatttt |
| KD246 | aaaataacccaaGAGAaaaGAGAaatGAGAaatacccaaa |
| KD247 | /5Cy5/tttTCTCattTCTCattgggttttTCTCttTCTCtatttt |
| KD248 | aaaataGAGAaaGAGAaaaacccaatGAGAaatGAGAaaa |
| T7 promoter with *Sa*Cas9 sgRNA insert for protein expression | gtcgacTAATACGACTCACTATAGGGTAATGAAATAAGATCACTACGTTTTAGTACTCTGGAAACAGAATCTACTAAAACAAGGCAAAATGCCGTGTTTATCTCGTCAACTTGTTGGCGAGATctcgag |
| KD153 | agatctcgagTGCGGCCGCACTCGAGCA |
| KD154 | attagtcgacAGCTTGTCGACGGAGCTCGAATTCG |
| KD155 | tcgacaagctGTCGACTAATACGACTCACTATAG |
| KD156 | tgcggccgcaCTCGAGATCTCGCCAACAAG |
| KD174 | tccttttcatGGATCCACCAATCTGTTCTCTGTGAGCCTCAATAATATC |
| KD175 | tggtggatccATGAAAAGGAACTACATTCTG |
| KD179 | agatctcgagGCGGCCGCACTCGAGGCC |
| KD180 | gtgcggccgcCTCGAGATCTCGCCAACAAG |
| *Sa*Cas9 SMART target sgRNA | UAAUGAAAUAAGAUCACUACGUUUUAGUACUCUGGAAACAGAAUCUACUAAAACAAGGCAAAAUGCCGUGUUUAUCUCGUCAACUUGUUGGCGAGAU |
| *Sa*Cas9 λ target-29.4 kb sgRNA | GCGAGGAUUGUUAUGUAAUAGUUUUAGUACUCUGGAAACAGAAUCUACUAAAACAAGGCAAAAUGCCGUGUUUAUCUCGUCAACUUGUUGGCGAGAU |
| IF365 | AAGAACGCCTCGCACACT |
| IF460 | /5atto647n/AACCGCCGAATAACAGAGT |
| *Sa*Cas9 SMART Target gBlock for non-target DNA EMSA | AAGAACGCCTCGCACACTcttttgacttgatcggcacgtaagaggttccaactttcaccataatgaaataagatcactacttgggttatttttgagttatcgagattttcagACTCTGTTATTCGGCGGTT |
| *Fn*Cas9 sgRNA gBlock for IVT | GGGATGTGCTGCAAGGCGATTAAGTTGGGTAACGCCAGGGTTTTCCCAGTCACGACGTTGTAAAACGACGGCCAGTGAGCGCGCGTAATACGACTCACTATAGGGataatgaaataagatcactacGTTTCAGTTGCTGAATTATTTGGTAAACAGTACCAAATAATTAATGCTCTGTAATCATTTAAAAGTATTTTGAACGGACCTCTGTTTGACACGTCTGAATAACTAAAAATTTTTTT |

| | |
|---|---|
| *Nme*Cas9 sgRNA gBlock for IVT | GGGATGTGCTGCAAGGCGATTAAGTTGGGTAACGCCAGGGTTTTCCCAGTCACGACGTTGTAAAACGACGGCCAGTGAGCGCGCGTAATACGACTCACTATAGGGaccataatgaaataagatcactacGTTGTAGCTCCCTTTCTCATTTCGGAAACGAAATGAGAACCGTTGCTACAATAAGGCCGTCTGAAAAGATGTGCCGCAACGCTCTGCCCCTTAAAGCTTCTGCTTTAAGGGGCATCGTTTA |
| KD142 | GGGATGTGCTGCAAGGCG |
| KD143 | TAAACGATGCCCCTTAAAGCAGA |
| KD144 | AAAAAAATTTTTAGTTATTCAGACGTGTCAAAC |
| ***Sa*Cas9 sgRNA for genome editing in HEK293T cells** | |
| KJ_T0217_CACNA1D_20nt_Fwd | CACCGGCAGGAGTATTTCAGTAGTG |
| KJ_T0221_CACNA1D_20nt_Rev | AAACCACTACTGAAATACTCCTGCC |
| KJ_T0324_EMX1_21nt_Fwd_3 | CACCGGGCCTCCCCAAAGCCTGGCCA |
| KJ_T0325_EMX1_21nt_Rev_3 | GAACTGGCCAGGCTTTGGGGAGGCCC |
| KJ_T0326_FANCF_21nt_Fwd_3 | CACCGGCAAGGCCCGGCGCACGGTGG |
| KJ_T0327_FANCF_21nt_Rev_3 | GAACCCACCGTGCGCCGGGCCTTGCC |
| KJ_T0328_RUNX1_23nt_Fwd_1 | CACCGGTACTCACCTCTCATGAAGCACT |
| KJ_T0329_RUNX1_23nt_Rev_1 | GAACAGTGCTTCATGAGAGGTGAGTACC |
| KJ_T0287_Scramble_sgRNA_Fwd | CACCGGTATTACTGATATTGGTGGG |
| KJ_T0288_scramble_sgRNA_Rev | GAACCCCACCAATATCAGTAATACC |
| **T7E1 PCR primers** | |
| KJ_T0225_CACNA1D_T7E1_Fwd | ACA GAC ACA CAC ACG GTG CT |
| KJ_T0226_CACNA1D_T7E1_Rev | TGG AGT TTC TGC TCC CAT TT |
| KJ_T0229_FANCF_T7E1_Fwd | ACC TCT TTG TGT GGC GAA AG |
| KJ_T0230_FANCF_T7E1_Rev | CCA GGC TCT CTT GGA GTG TC |
| KJ_T0231_EMX1_T7E1_Set1_Fwd | GCC CCT AAC CCT ATG TAG CC |
| KJ_T0232_EMX1_T7E1_Set1_Rev | GGA GAT TGG AGA CAC GGA GA |
| KJ_T0320_RUNX1_PCR_Fwd | CCAGCACAACTTACTCGCACTTGAC |

| | |
|---|---|
| KJ_T0321_RUNX1_PCR_Rev | CATCACCAACCCACAGCCAAGG |
| KJ_T0206_AcrIIA11a1 gBlock for expression in HEK293T cells | tataggGagacccaagctggctagcATG GCA GAT ATG ACG CTT CGC CAG TTC TGCGAG CGA TAT CGC AAG GGT GAC TTC CTC GCAAAG GAT CGA GAA ACT CAA ATC GAG GCA GGTTG G TAC GAT TGG TTT TGT GAT GAC AAA GCCTTG GCG GG C CGA TTG GCA AAA ATC TGG GGGATT TTG AAG GGG A TA ACC TCA GAT TAT ATCTTG GAT AAC TAC CGC GTA T GG TTC AAA AACAAC TGT CCA ATG GTA GGA CCA CTG TAC GACGAT GTA CGC TTC GAA CCG CTT GAT GAA GAA CAG CGA GAT GAG CTC TAC TTC GGC GTC GCAATC GAC GAT AAG AGG AGG GAA AAG AAA TACGTC ATA TTC AC T GCT CGA AAT GAC TAT GAAAAC GAG TGT GGT TTC A AC AAC GTG AGA GAAGTA CGC CAA TTT ATA AAT GGA TGG GAA GACGAA TTG AAG AAC GAA GAG TTC TAT AA G GCTAGG GAG AAA AAA CGG CAA GAA ATG GAA GAA GCC AAT AAC AAA TTC GCA GAA ATA ATG CAACGG GC C GAT GAG ATA TTG TGG AAC CTG AAAGAG GACtccggacc tccgaagaaaaagcgaaaggtg ggatccagtgga tatccctatgacgtgcccgattatgcc taaCtcgagcggccgccactgtgctgga |

**Table S2. Plasmids used in this study.**

| Plasmid | Description | Primers | Source |
|---|---|---|---|
| pIF592 | p6XHis_NLS-*Sa*Cas9 (item #101086) | N/A | (Soares et al., 2017) |
| pIF936 | pMCSG7-Wt-*Nme*Cas9 (item # 71474) | N/A | (Zhang et al., 2015) |
| pIF937 | AcrIIA11-6xHis in pET19 vector | N/A | This study |
| pIF938 | AcrIIA11-TS in pET19 vector | N/A | This study |
| pIF939 | TS-SUMO-AcrIIA11 No C-terminal tags in pET19 vector | N/A | This study |
| pIF940 | TS-SUMO-3xFLAG-*Sa*Cas9 | N/A | This study |
| pIF941 | TS-SUMO-3xFLAG-d*Sa*Cas9 | KD197 and KD198 | This study |
| pIF942 | TS-SUMO-*Sa*Cas9-sgRNA | KD174, KD175, KD179, KD180 | This study |
| pIF943 | p6xHis_NLS-*Sa*Cas9-sgRNA | KD153, KD154, KD155, KD156, T7 promoter with *Sa*Cas9 sgRNA insert | This study |

| | | | |
|---|---|---|---|
| pIF944 | pCK002_U6-Sa-sgRNA(mod)_EFS-SaCas9-2A-Puro_WPRE (item # 85452) | KJ_T0217_CACNA1D_20nt_Fwd, KJ_T0221_CACNA1D_20nt_Rev, KJ_T0324_EMX1_21nt_Fwd_3, KJ_T0325_EMX1_21nt_Rev_3, KJ_T0326_FANCF_21nt_Fwd_3, KJ_T0327_FANCF_21nt_Rev_3, KJ_T0328_RUNX1_23nt_Fwd_1, KJ_T0329_RUNX1_23nt_Rev_1, KJ_T0287_Scramble_sgRNA_Fwd, KJ_T0288_scramble_sgRNA_Rev, | (Singer et al., 2016) |
| pIF945 | pAAV-CMV-NLS-AcrIIA4 (item # 113038) | N/A | (Bubeck et al., 2018) |
| pIF946 | pAAV-CMV-NLS-AcrIIA11 | N/A | This study |
| pIF967 | TS-SUMO-*Sa*Cas9 | N/A | This study |

**Table S3. Single-molecule data analysis.**

| **Diffusing vs stationary *Sa*Cas9 molecules (Figure 3C)** | | | | |
|---|---|---|---|---|
| Condition | Stationary molecules | Diffusing molecules | Number of molecules | p-value (Chi-squared test) |
| - AcrIIA11 | 28 | 61 | 89 | $3 \times 10^{-9}$ |
| + AcrIIA11 | 70 | 23 | 93 | |
| ***Sa*Cas9 diffusion coefficients (Figure 3E)** | | | | |
| Condition | Mean diffusion coefficient ± S.E.M. ($\mu m^2$ $s^{-1}$) | | Number of molecules | p-value (Mann-Whitney U-test) |
| - AcrIIA11 | 0.05 ± 0.01 | | 33 | $9.6 \times 10^{-7}$ |
| + AcrIIA11 | 0.006 ± 0.003 | | 33 | |
| **WT *Sa*Cas9 target binding (Figure 4D)** | | | | |
| Condition | Target bound molecules | Non-target bound molecules | Number of molecules | p-value (Chi-squared test) |
| - AcrIIA11 | 38 | 51 | 89 | $1 \times 10^{-6}$ |
| + AcrIIA11 | 13 | 95 | 108 | |
